# Fecal microbiota transplantation mitigates cardiac remodeling and functional impairment in mice with chronic colitis

**DOI:** 10.1101/2025.03.13.643179

**Authors:** Xiaoying S Zhong, Kevin M. Lopez, Srikruthi S. Krishnachaitanya, Max Liu, Ying Xiao, Rongliwen Ou, Hania I. Nagy, Thierry Kochkarian, Don W. Powell, Ken Fujise, Qingjie Li

## Abstract

**Background:** Inflammatory bowel disease (IBD) is a chronic inflammatory disorder with significant extraintestinal manifestations, including cardiovascular derangements. However, the molecular mechanisms underlying the cardiac remodeling and dysfunction remain unclear.

**Methods:** We investigated the effects of chronic colitis on the heart using two mouse models: DSS-induced colitis and *Il10^-/-^*spontaneous colitis. Echocardiography was employed to assess heart function and molecular characterization was performed using bulk RNA-sequencing, RT-qPCR, and western blot.

**Results:** Both models exhibited significant cardiac impairment, including reduced ejection fraction and fractional shortening as well as increased collagen deposition, inflammation, and myofibril reorganization. Molecular analyses revealed upregulation of fibrosis markers (i.e. COL1A1, COL3A1, Fibronectin) and β-catenin reactivation, indicating a pro-fibrotic cardiac environment. Each model yielded common upregulation of eicosanoid-associated and inflammatory genes (*Cyp2e1*, *Map3k6*, *Pck1*, *Cfd*), and model-specific alterations in pathways regulating cAMP- and cGMP-signaling, arachidonic and linoleic acid metabolism, Cushing syndrome-related genes, and immune cell responses. DSS colitis caused differential regulation of 232 cardiac genes, while *Il10^-/-^* colitis yielded 105 dysregulated genes, revealing distinct molecular pathways driving cardiac dysfunction. Importantly, therapeutic fecal microbiota transplantation (FMT) restored heart function in both models, characterized by reduced fibrosis markers and downregulated pro-inflammatory genes (*Lbp* and *Cdkn1a* in *Il10^-/-^* mice and *Fos* in DSS mice), while also mitigating intestinal inflammation. Post-FMT cardiac RNA-sequencing revealed significant gene expression changes, with three altered genes in DSS mice and 67 genes in *Il10^-/-^* mice. Notably, *Il10^-/-^* mice showed relatively less cardiac recovery following FMT, highlighting IL-10’s cardioprotective and anti-inflammatory contribution.

**Conclusions:** Our findings elucidate novel insights into colitis-induced cardiac remodeling and dysfunction and suggest that FMT mitigates cardiac dysfunction by attenuating systemic inflammation and correcting gut dysbiosis. This study underscores the need for further evaluation of gut-heart interactions and microbiome-based therapies to improve cardiovascular health in IBD patients.

**Graphical Abstract:** 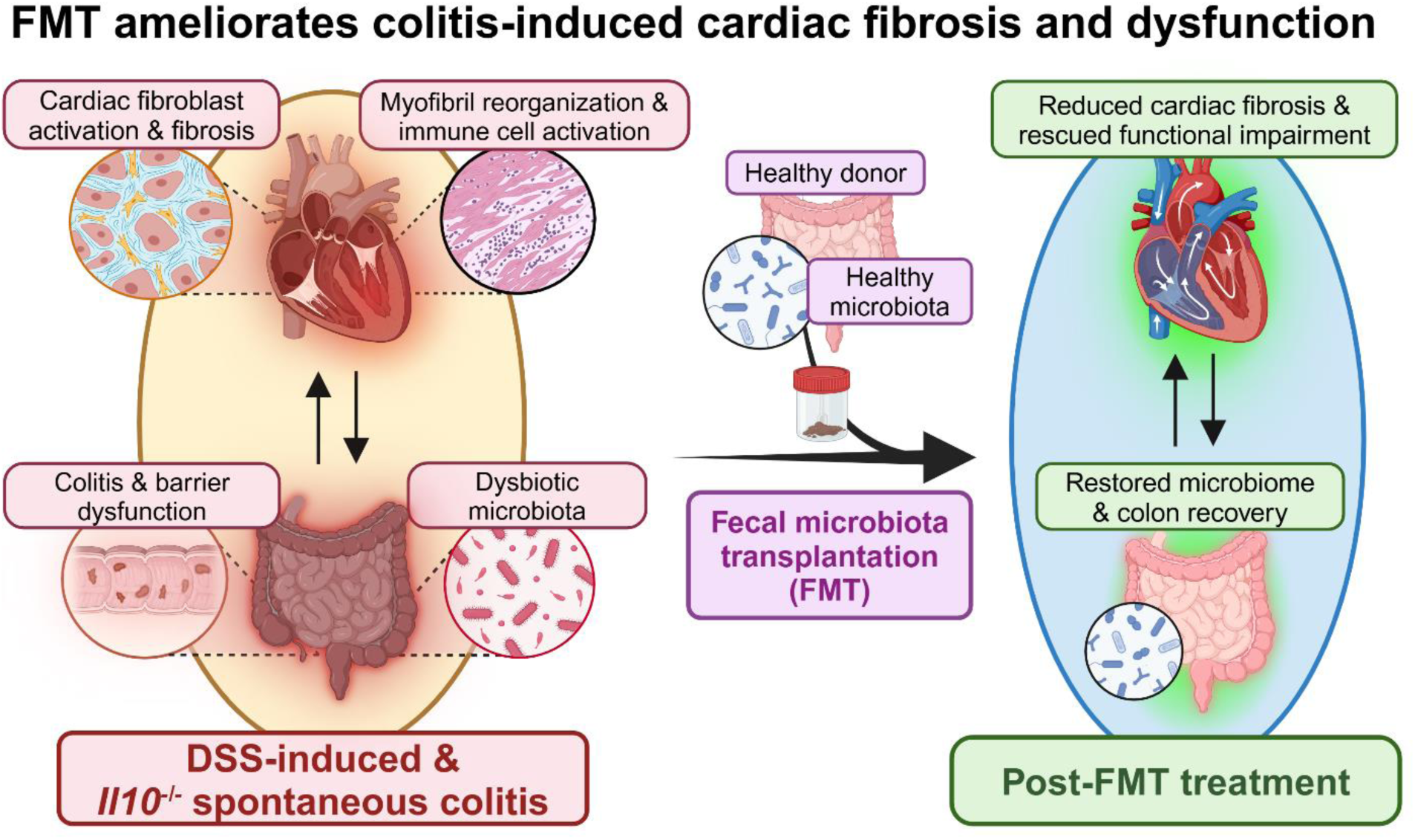

## 1. Introduction

Inflammatory bowel diseases (IBDs) are chronic conditions characterized by prolonged, recurrent inflammation of the gastrointestinal tract, varying in anatomic location, histopathologic morphology, and molecular pathophysiology, often rendering patients debilitated by its effects. The two most common IBDs are ulcerative colitis and Crohn’s disease, which are known to present with various extraintestinal manifestations and complications such as in the skin (e.g. erythema nodosum), eyes (e.g. uveitis), joints/bones (e.g. enteropathic arthritis), blood (e.g. iron-deficiency anemia), hepatobiliary system (e.g. primary sclerosing cholangitis), and urogenital system (e.g. urolithiasis) (1). Recently, research has elucidated a relationship between IBD and cardiovascular diseases in the Gut-Heart-Axis. Cardiac manifestations of IBD such as cardiomyopathy, arrythmia, myocardial infarction (MI), pericarditis, and myocarditis have been documented. Various cohort studies evaluating cardiac manifestations in IBD patients report increased risk of acute MI, atrial fibrillation, pericarditis or myocarditis, and an overall increased risk of hospitalization for heart failure (2–4). Additionally, vascular manifestations of IBD have been reported such as atherosclerosis and coronary artery disease, arterial/venous thromboembolism, and vascular remodeling (2, 5–7). Further understanding of the molecular mechanisms by which cardiovascular manifestations of IBD occur becomes important, highlighting a need for further basic science research.

Recently, we determined that DSS-induced chronic colitis causes an interleukin-1 beta (IL-1β)-mediated increase in miR-155, yielding brain-derived neurotrophic factor (BDNF) suppression, reduced left ventricular ejection fraction (LVEF) with increased end systolic volume, and left ventricular hypertrophy, demonstrating molecular cardiac reorganization and functional impairment. Treatment of IBD using anti-IL-1β antibody mitigated DSS-induced cardiac remodeling via miR-155/BDNF signaling (8). Furthermore, we demonstrated that an increased gut-derived exosomal miR-29b induces cardiac remodeling and impairment via suppression of important cardiac proteins, especially BDNF (9). However, greater understanding of pathogenic cardiac remodeling due to IBD remains important, paving the way for discovery of novel therapeutic agents to treat these cardiac complications.

Therapeutic interventions for IBD focus on induction of remission with long-term asymptomatic maintenance. Current pharmacologic therapies utilize anti-inflammatory agents including corticosteroids (e.g. budesonide), biologics (e.g. anti-tumor necrosis factor alpha antibodies), immunomodulators (e.g. thiopurine analogs), and 5-aminosalicyclic acid derivatives (5-ASA) (e.g. sulfasalazine) (10, 11). Newer pharmacotherapies with promising outcomes in clinical practice include sphingosine-1-receptor modulators such as etrasimod and ozanimod (12, 13). Historically, extraintestinal manifestations of IBD are treated according to respective symptoms, but the underlying pathophysiology and most effective therapeutic intervention for cardiac manifestations of IBD must still be elucidated. The aim of this study is to clarify the impact that chronic colitis has on cardiac function and molecular programming using two independent mouse models and how restoring the gut microbiota symbiosis using therapeutic fecal microbiota transplantation (FMT) may repair defects in cardiac function and histology. Our results demonstrate that each chronic colitis model induces significant reductions in functional parameters of the heart accompanied by marked alterations to the cardiac transcriptome and increased expression of fibrotic and profibrotic proteins. Therapeutic FMT ameliorated IBD-induced cardiac dysfunction and rescued some dysregulated cardiac genes in each model, though, *Il10^-/-^* mice demonstrated relatively less recovery compared to DSS mice. Importantly, FMT rescued cardiac dysfunction in the absence of IL-10, suggesting that the gut microbiota play a unique role in development of cardiovascular defects during chronic colitis, which FMT can selectively target while simultaneously improving intestinal symptoms. Our findings underscore the intricate link between the gut microbiome and heart health, highlight the importance of long-term monitoring of heart function in IBD patients, and suggest a potential therapeutic role for FMT in managing cardiovascular manifestations associated with IBD.

## 2. Methods

### 2.1. Reagents

Dextran sodium sulfate (DSS), molecular weight 36-50K, was purchased from Cayman Chemical (Ann Harbor, MI).

### 2.2. Animals

Six-week-old male C57BL/6J mice were purchased from the Jackson Laboratory (Bar Harbor, ME) and housed in the UTMB Animal Facility for two weeks to allow for acclimation. The animals were randomized into 3 groups of 10 mice each: control group, DSS group, and DSS+FMT group. Mice from the control group served as fecal donors for FMT. One day before the DSS treatment, fecal pellets were collected, weighted, and dispersed in PBS containing 15% glycerol (14). The fecal slurry was filtered, aliquoted, and stored in a -80°C freezer. Sterilized fecal slurry was included as vehicle control. To induce chronic colitis, mice were given 2% DSS in drinking water for 4 cycles of 7 days each, with intermittent 2-week intervals of normal drinking water. On the last day of each DSS cycle, 200 µL of fecal slurry or vehicle control was administered via oral gavage. Mice were euthanized 2 weeks after the last FMT. The procedures were approved by the IACUC at the University of Texas Medical Branch (IACUC # 1512071B).

Heterozygous *Il10*^-/+^ mice on the 129/Sv background (129(B6)-*Il10^tm1Cgn^*/J, strain # 004368) were also purchased from the Jackson Laboratory. After breeding, homozygous *Il10*^-/-^ mice were utilized for the experiments and homozygous *Il10*^+/+^ mice served as controls. Mice were housed under specific pathogen-free (SPF) conditions in the UTMB Animal Facility with a 12-hour light/dark cycle, a temperature range of 24-26°C, and a relative humidity of 40-70%. When the animals were 2 months old, fecal pellets were collected from the *Il10*^+/+^ mice and processed as described above. Mice in the *Il10*^-/-^+FMT group were treated with 200 µL of fecal slurry via oral gavage monthly for 8 months and mice in the *Il10*^-/-^ group received sterilized fecal slurry at the same interval. All *Il10*^+/+^ and *Il10*^-/-^ mice were euthanized at 10 months of age.

All mice were euthanized under a deep plane of anesthesia with isoflurane. The individual heart was removed, rinsed in PBS, blotted dry, and cut into two halves, one snap-frozen in liquid nitrogen for molecular analyses and another fixed in formalin for histology. The whole colons were rinsed with ice-cold saline, opened longitudinally, snap-frozen in liquid nitrogen or pinned flat, Swiss-rolled, and fixed in 10% neutral buffered formalin for histological analysis.

### 2.3. Mouse echocardiography

Mouse cardiac function was evaluated via trans-thoracic echocardiography (TTE) using a Vevo 770 High-Resolution Imaging System (VisualSonics, Toronto, Canada) (8). Mice were anesthetized with isoflurane (4% for induction and 1% for maintenance) and positioned in left lateral decubitus. The anterior chest hair was removed with Nair Hair Remover Lotion (Church & Dwight Co., Ewing, NJ). ECG was monitored throughout the experiment and temperature was kept at 37°C on a heated plate. The left ventricular short-axis and long-axis view and the apical four-chamber view were examined by two-dimensional and M-mode echocardiography. Cardiac function-related parameters were determined, including LVEF and fractional shortening (FS).

### 2.4. Hematoxylin and eosin (H&E) staining

Mouse heart tissue was fixed in formalin for 48 hours and embedded in paraffin. For histologic examination, five-micrometer tissue sections were stained with H&E (15). Images were captured using a LEICA DMI 6000 B microscope.

### 2.5. Masson’s trichrome staining

Collagenous connective tissue fibers in heart sections were stained using a Trichrome stain kit (Cat. # ab150686. Abcam, Cambridge, UK).

### 2.6. Myeloperoxidase (MPO) assay

Frozen colons were pulverized in liquid nitrogen. A predetermined aliquot of tissue powder was homogenized in 20 mM phosphate buffer (pH 7.4) and centrifuged at 4°C for 10 minutes (16). The pellets were then sonicated in 50 mM phosphate buffer (pH 6.0) containing 0.5% hexadecyl trimethyl ammonium bromide and centrifuged at 4°C for 5 minutes. The supernatant (100 μL) was collected and incubated with 16 mM tetramethyl benzidine in 50% ethanol, 0.3 mM H_2_O_2_, and 8 mM sodium phosphate buffer (pH 5.4) for 3 minutes. The optical density (OD) was measured at 655 nm.

### 2.7. Bulk RNA-sequencing (RNA-seq)

Total RNAs were isolated from pulverized frozen heart tissue using the PureLink RNA Mini Kit (ThermoFisher Scientific, Waltham, MA) (17). Libraries were prepared via the Illumina TruSeq RNA Sample Preparation kit (18). Cluster formation of the library DNA templates was performed using the TruSeq PE Cluster Kit v3 and the Illumina cBot workstation. Paired-end 50 base sequencing by synthesis was performed using TruSeq SBS kit v3 on an Illumina HiSeq 1000. The alignment of sequence reads was performed using the Spliced Transcript Alignment to a Reference program version 2.4.2a, using default parameters. Differential expression was evaluated using Cufflinks/Cuffdiff software, v2.2.1. Gene ontology (GO) and Kyoto Encyclopedia of Genes and Genomes (KEGG) pathway enrichment within the gene lists of interest were performed using Enrichr (https://maayanlab.cloud/Enrichr/).

### 2.8. Reverse transcriptase-quantitative polymerase chain reaction (RT-qPCR)

Complementary DNAs were synthesized using the M-MuLV Reverse Transcriptase (NEB, Ipswich, MA). qPCR was performed using the PowerUp SYBR Green Master Mix (ThermoFisher) (15). Glyceraldehyde 3-phosphate dehydrogenase (*Gapdh*) served as internal control. Primer sequences are listed in Table S1.

### 2.9. Western blot (WB)

To detect protein expression levels in the myocardium, WB was performed as described previously (16). Primary antibodies are listed in Table S2.

### 2.10. Statistical analysis

The comparisons of means among groups were analyzed by t-test or one way ANOVA, and the Dunn Multiple Comparison Test was further used to determine significant differences between groups. All statistical analyses were performed using the SPSS package (version 13.0). A value of *p<*0.05 was considered statistically significant.

### 2.11. Data Availability

Data supporting this study’s findings are available from the corresponding author upon reasonable request.

## 3. Results

### 3.1. Chronic colitis was ameliorated by FMT in DSS-treated and Il10^-/-^ mice

The detrimental impact of IBD on overall health has been well established in humans hallmarked by dysbiosis, inflammation, and weight loss due to anorexia and malabsorption (19) and animal models are essential for mechanistic and therapeutic evaluation. To determine the impact of chronic colitis on heart function and cardiac remodeling, we chose two independent mouse models: 1) DSS-induced chronic colitis model and 2) *Il10^-/-^ mice with* spontaneous colitis (20, 21). Here, we first verified colonic inflammation and evaluated the impact of FMT on their inflammatory states. DSS-treated mice showed significantly reduced body weight and the reduction was significantly mitigated by FMT (Figure 1A). Body weights of *Il10^-/-^* mice were also significantly reduced compared to wild type (WT) mice while FMT showed only marginal recovery in body weight (Figure 1B). Colon lengths of DSS-treated mice were significantly shortened compared to controls and FMT significantly mitigated the colonic shortening (Figure 1C). The colon lengths of *Il10^-/-^* mice are similar to that of WT mice but FMT had little effect (Figure 1D). RT-qPCR detected significantly higher levels of interleukin 1 beta (*Il1b)* mRNA expression in the colons of DSS-treated mice while FMT ameliorated this increase (Figure 1E). Similarly, *Il1b* mRNA overexpression was observed in *Il10^-/-^*mice and the elevation was significantly ameliorated by FMT (Figure 1F). Additionally, MPO activity was shown to be significantly upregulated in DSS mouse colons, further mitigated by FMT (Figure 1G). Coincidingly, *Il10^-/-^* mice showed significantly greater MPO activities where FMT effectively reduced the increases (Figure 1H). Overall, chronic inflammation was verified in the colons of C57BL/6J mice subjected to 4 intermittent DSS insults and 10 months old *Il10^-/-^* mice. Therapeutic FMT treatment was effective at mitigating, but not eradicating, chronic colitis in both DSS and *Il10^-/-^* mice.

**Fig. 1.**
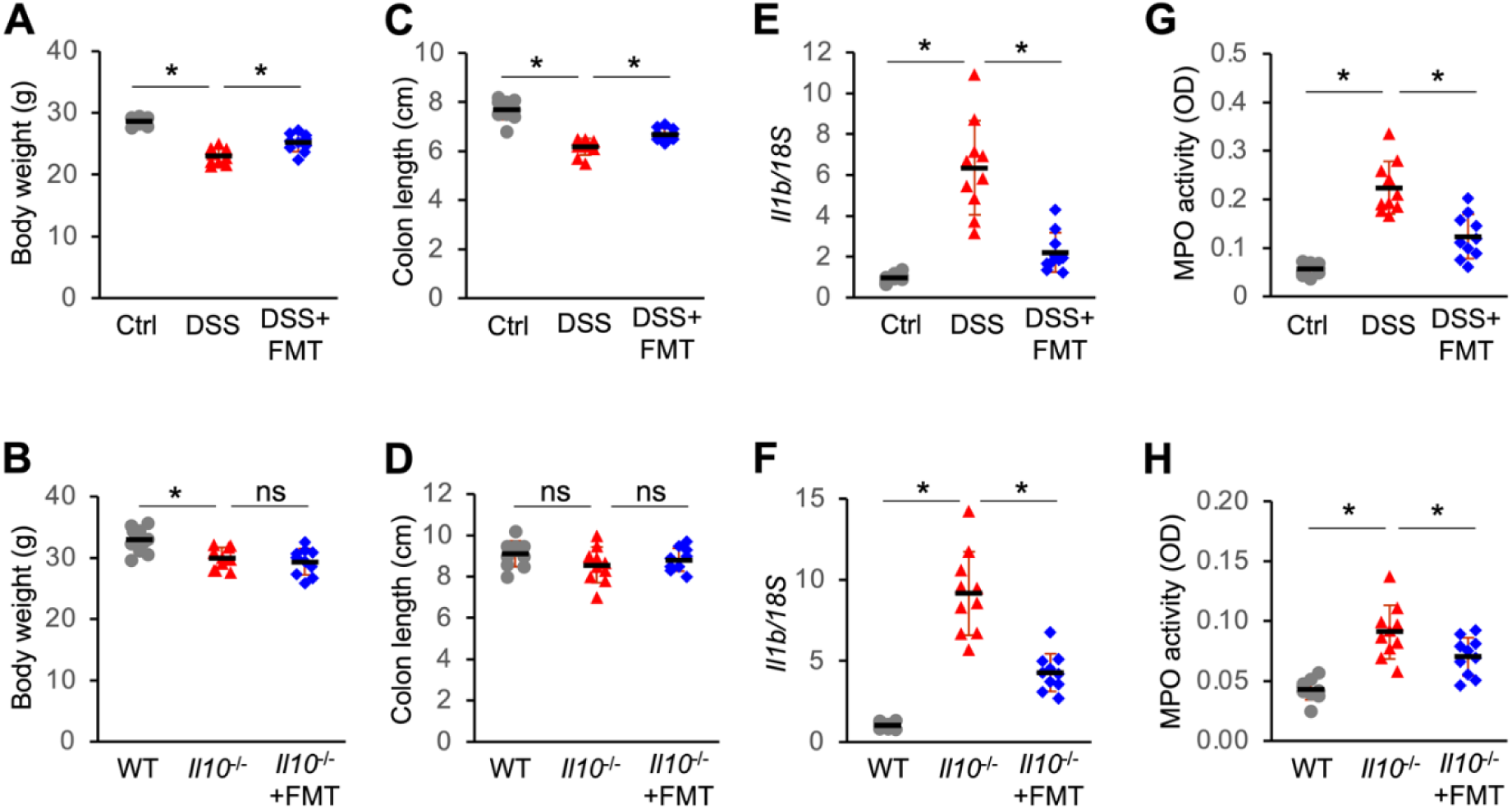
Fecal microbiota transplantation (FMT) significantly mitigated colonic inflammation in DSS-treated and *Il10*^-/-^ mice. (***A***) Body weights of DSS mice treated with or without FMT. (***B***) Body weights of wild type mice, *Il10*^-/-^ mice, and *Il10*^-/-^ mice subjected to FMT. (***C***) DSS-induced colon length shortening was ameliorated by FMT. (***D***) Spontaneous colitis and FMT had minimal effect on colon lengths in *Il10*^-/-^ mice. (***E***) DSS-induced upregulation in *Il1b* mRNA was mitigated by FMT. (***F***) FMT rescued *Il1b* mRNA overexpression in *Il10*^-/-^ mice. (***G***) DSS markedly increased myeloperoxidase (MPO) activities in the colon and the increase was mitigated by FMT. (***H***) The increase of MPO activities in *Il10*^-/-^ mice was also significantly ameliorated by FMT. n=10. One-way ANOVA. * *p*<0.05. ns, not significant.

### 3.2. Heart function impairment was mitigated by FMT in both DSS and Il10^-/-^ mice

Next, we evaluated heart function in DSS and *Il10^-/-^* mice and their respective controls using TTE. Chronic DSS colitis significantly reduced LVEF (Figure 2A) and FS (Figure 2B), indicative of heart function impairment. Of note, FMT effectively ameliorated these reductions (Figure 2A and B). H&E staining revealed increased inflammatory cell infiltration, myofibril disorganization, and interstitial edema in DSS mice compared to the controls; these changes were ameliorated by FMT treatment (Figure 2C). In *Il10^-/-^* mice, significantly less LVEF (Figure 2D) and FS (Figure 2F) were observed when compared to WT mice, with FMT mitigating these decreases like in DSS mice (Figure 2D and 2E). Compared to WT mice, *Il10^-/-^* mice had greater inflammatory cell infiltration, myofibril disorganization, and interstitial edema in the hearts and the increases were abrogated by FMT (Figure 2F). These results highlight the cardiac impairment in the presence of chronic colitis, as well as the therapeutic potential of FMT in improving cardiac functional parameters in patients with IBD.

**Fig. 2.**
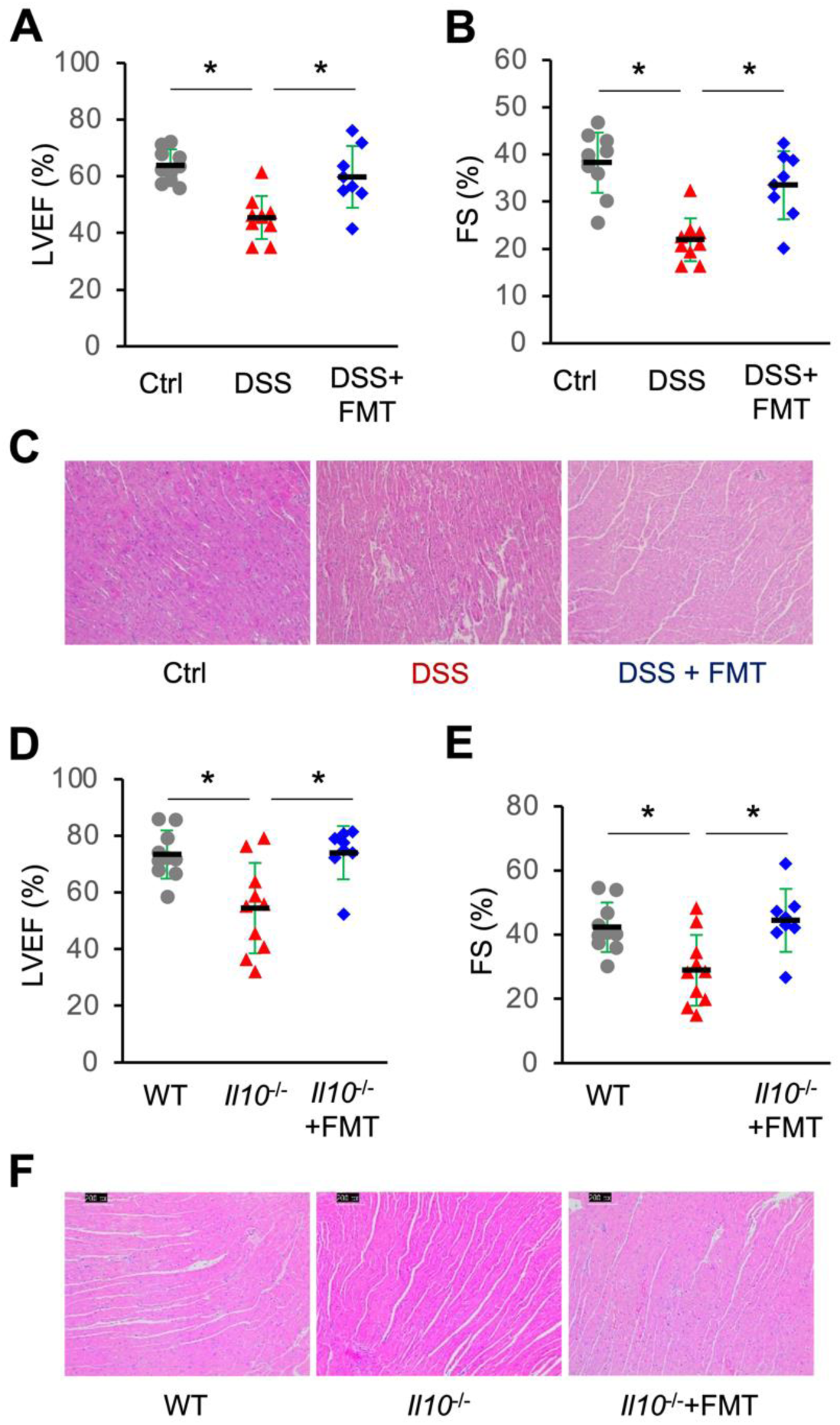
FMT significantly ameliorated colitis-induced cardiac impairment. Repeat DSS treatments significantly decreased left ventricular ejection fraction (LVEF, ***A***) and fractional shortening (FS, ***B***) and the decreases were mitigated by FMT. (***C***) H&E staining of the heart sections from control (Ctrl), DSS, and DSS+FMT mice (10X magnification). Spontaneous colitis also significantly suppressed LVEF (***D***) and FS (***E***) in 10 months old *Il10*^-/-^ mice and the suppression was ameliorated by FMT. n=10. One-way ANOVA. \**p*<0.05. (***F***) H&E staining of the cardiac sections from wild type, *Il10*^-/-^, and *Il10*^-/-^+FMT mice (10X magnification).

### 3.3. DSS colitis significantly altered the cardiac transcriptome profile

To further understand the mechanisms underlying the observed reduction in heart functionality, we profiled cardiac transcriptomes using bulk RNA-seq and identified signaling pathways involved in the process. Principal component analysis (PCA) of DSS heart transcriptome demonstrated clustering of control groups and DSS-treatment groups, respectively (Figure 3A), indicating a significant difference between the two groups. Further analysis of differential expression of heart mRNAs between DSS-treated and control mice revealed 232 significantly dysregulated genes of which 137 genes were upregulated and 95 genes were downregulated compared to controls (Figure 3B and Table S3). Among the 137 upregulated genes, the top 10 were *Gm6969*, *Ppm1k*, *Gdpd3*, *Gm3375*, *Cirbp*, *Adam19*, *Osbpl3*, *Tmprss13*, *Tceal8*, and *Elovl7*. Among the 95 downregulated genes, the top 10 were *Myot*, *Gm43305*, *Ano5*, *St3gal5*, *Zfp970*, *Serpine1*, *Tnfrsf12a*, *Lyve1*, *Mt2*, and *mt-Ti*. Hierarchal cluster analysis unveiled the top 30 differentially expressed genes (DEGs) between DSS-treated and control mice (Figure 3C) and the top 20 differentially expressed cardiac genes are presented in Table 1. KEGG pathway analysis of the 232 DEGs revealed 23 enriched terms with statistical significance (Table S4). The top 10 enriched signaling pathways include oxytocin regulation pathway, dopaminergic synapse, apelin signaling pathway, PPAR signaling pathway, cAMP signaling pathway in addition to pathways associated with Cushing syndrome, cocaine addiction, regulation of lipolysis in adipocytes, circadian entrainment, and parathyroid hormone synthesis, secretion, and action. GO enrichment analysis identified 102 dysregulated biological processes (Table S5). The top 10 biological processes include regulation of metal ion transport, regulation of sodium ion transport, negative regulation of synaptic transmission, positive regulation of tau-protein kinase activity, regulation of cGMP-mediated signaling, positive regulation of metabolic process in addition to ketone body biosynthetic process, kidney development, ketone body metabolic process, and response to alkaloid.

**Fig. 3.**
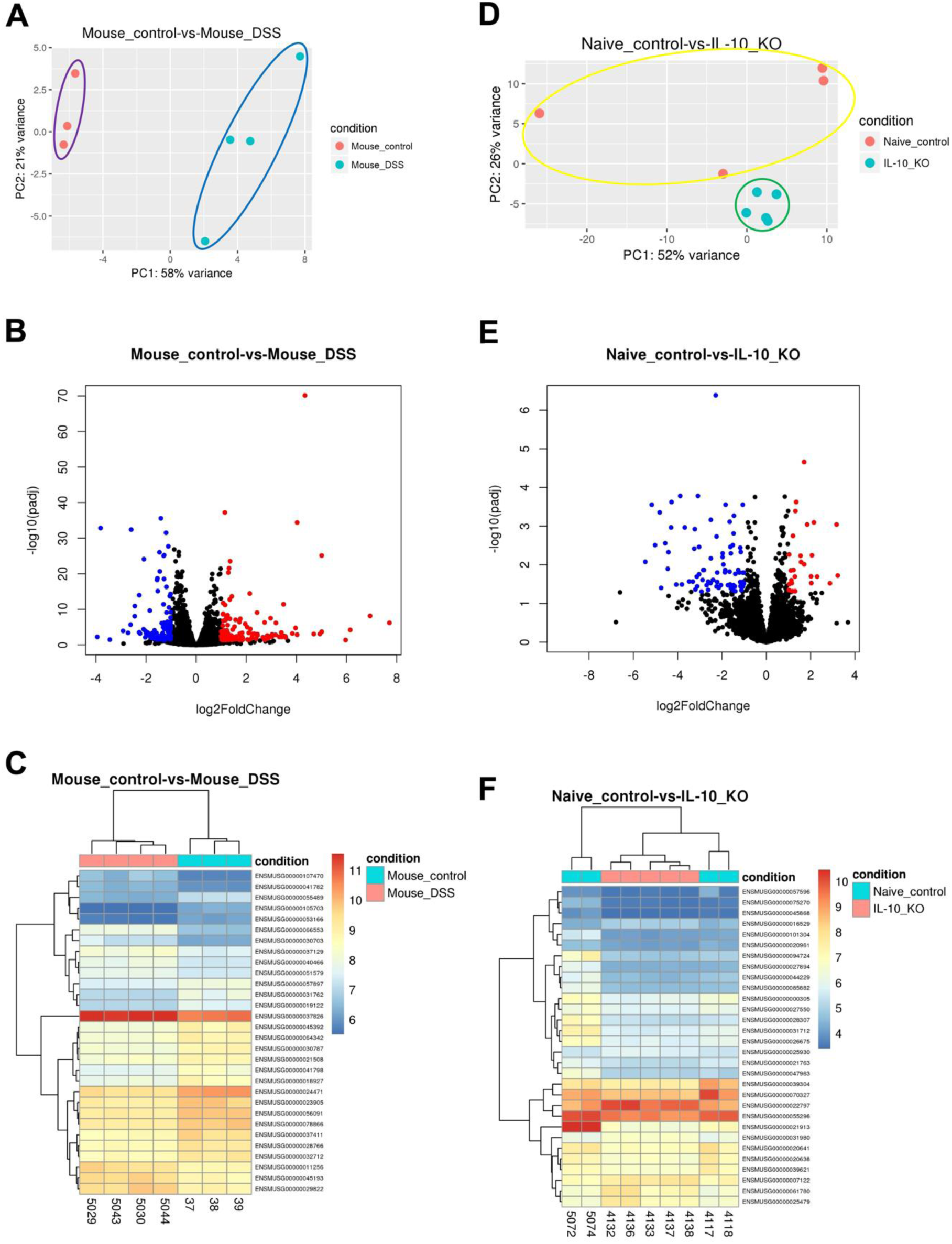
Bulk RNA-sequencing analysis of cardiac transcriptomes in DSS-treated and *Il10*^-/-^ mice. (***A***) Principal component analysis (PCA) of DSS mice (orange circles) vs control mice (turquoise circles). (***B***) Volcano plot representation of differential expression analysis of cardiac mRNAs in DSS mice vs control mice. Blue circles represent downregulated genes, and red circles represent upregulated genes. (***C***) Hierarchical clustering of normalized mRNA levels for the top 30 mRNAs having statistical significance for differential expression between DSS and control mice. (***D***) PCA of *Il10*^-/-^ vs wild type mice. (***E***) Volcano plot representation of differential expression analysis of cardiac mRNAs in *Il10*^-/-^ vs wild type mice. (***F***) Hierarchical clustering of normalized mRNA levels for the top 30 mRNAs having statistical significance for differential expression between *Il10*^-/-^ and wild type mice. n=3 or 4.

**Table 1.**
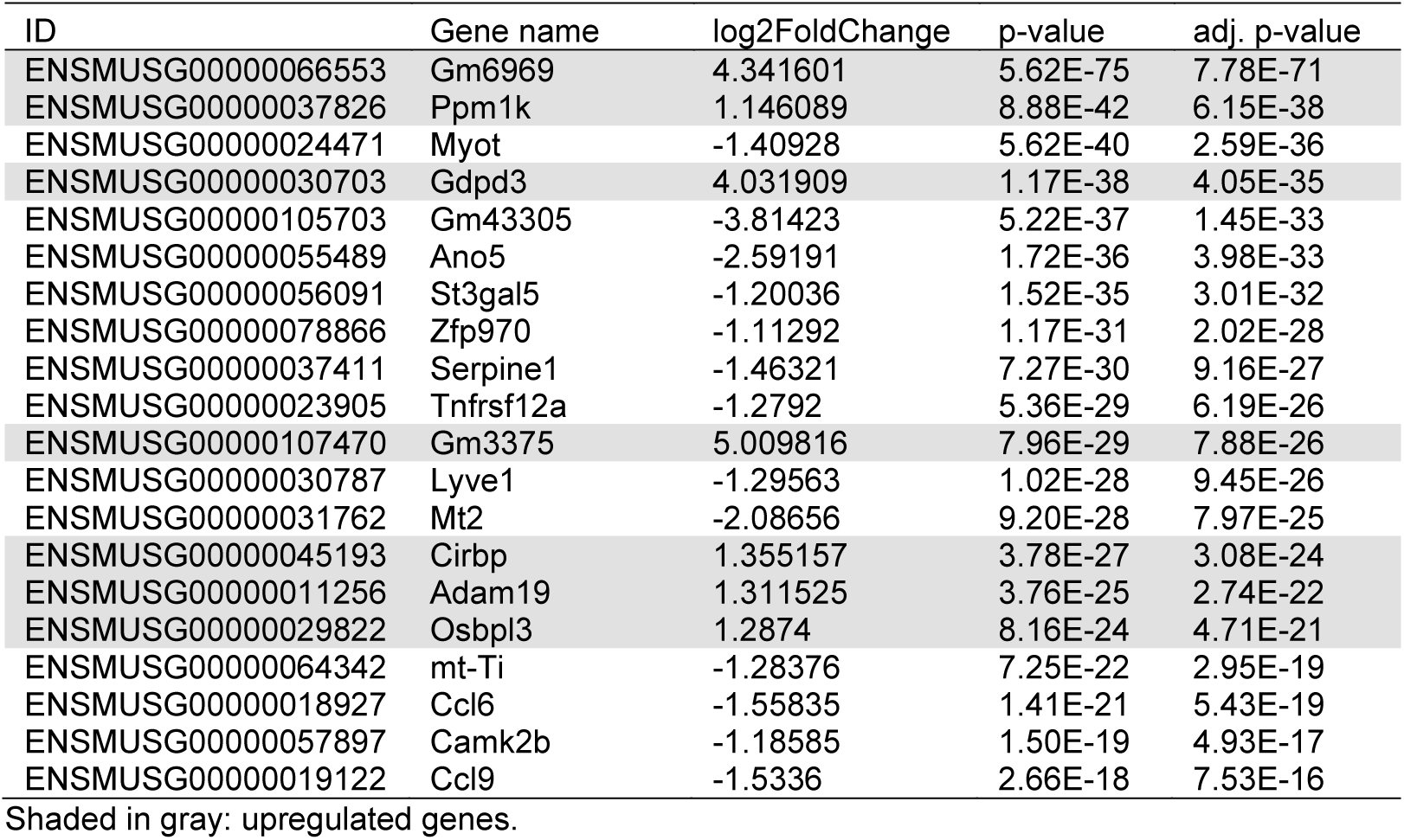
Top 20 cardiac genes significantly altered by DSS in C57BL/6 mice.

### 3.4. Spontaneous colitis also induced significant alterations in cardiac transcriptome in Il10^-/-^ mice

We also evaluated the impact of spontaneous colitis on the cardiac transcriptome in 10-month-old *Il10^-/-^* mice. PCA revealed a clustering of *Il10^-/-^* mice independent of *Il10^+/+^* WT mice. However, *Il10^+/+^* mice didn’t cluster well, demonstrating some variation among controls (Figure 3D). Volcano plot analysis identified 105 significantly dysregulated genes in *Il10^-/-^* mice, of which 26 genes were upregulated, and 79 genes were downregulated compared to controls (Figure 3E and Table S6). We report the top 20 dysregulated genes with the highest valued statistical significance in *Il10^-/-^* mice (Table 2). Hierarchal clustering of the top 30 DEGs is also presented (Figure 3F). The top 10 significantly upregulated genes were *Tfrc*, *Casq1*, *Agt*, *Cfd*, *Cyp2e1*, *Msc*, *Pla2g4e*, *Cidec*, *Shisha2*, and *Retnla*, while the top 10 downregulated genes were *Cdh4*, *Il15*, *Nxpe4*, *Slc6a17*, *BC067074*, *Ogdhl*, *Tmem245*, *Plet1os*, *Cmpk2*, and *Ston2*. KEGG pathway analysis of the 105 DEGs revealed 15 enriched terms with significant p-values (Figure S7). The top 10 enrichments include linoleic acid metabolism, citrate cycle, NOD-like receptor signaling pathway, insulin resistance, intestinal immune network for IgA production, ovarian steroidogenesis, FoxO signaling pathway, vascular smooth muscle contraction, steroid hormone biosynthesis, and arachidonic acid metabolism. GO biological process analysis of the 105 DEGs revealed enrichments in 120 biological processes (Table S8). The top 10 enriched terms include positive regulation of immune response, regulation of T cell proliferation, regulation of interleukin-18 production, defense response to symbiont, granulocyte activation, regulation of isotype switching, defense response to virus, neutrophil activation, positive regulation of T cell proliferation, and positive regulation of lymphocyte proliferation. The transcriptomic and metabolic changes observed in the hearts of each mouse model highlights the fundamental underlying molecular reprogramming occurring in cardiac tissue during chronic colitis.

**Table 2.**
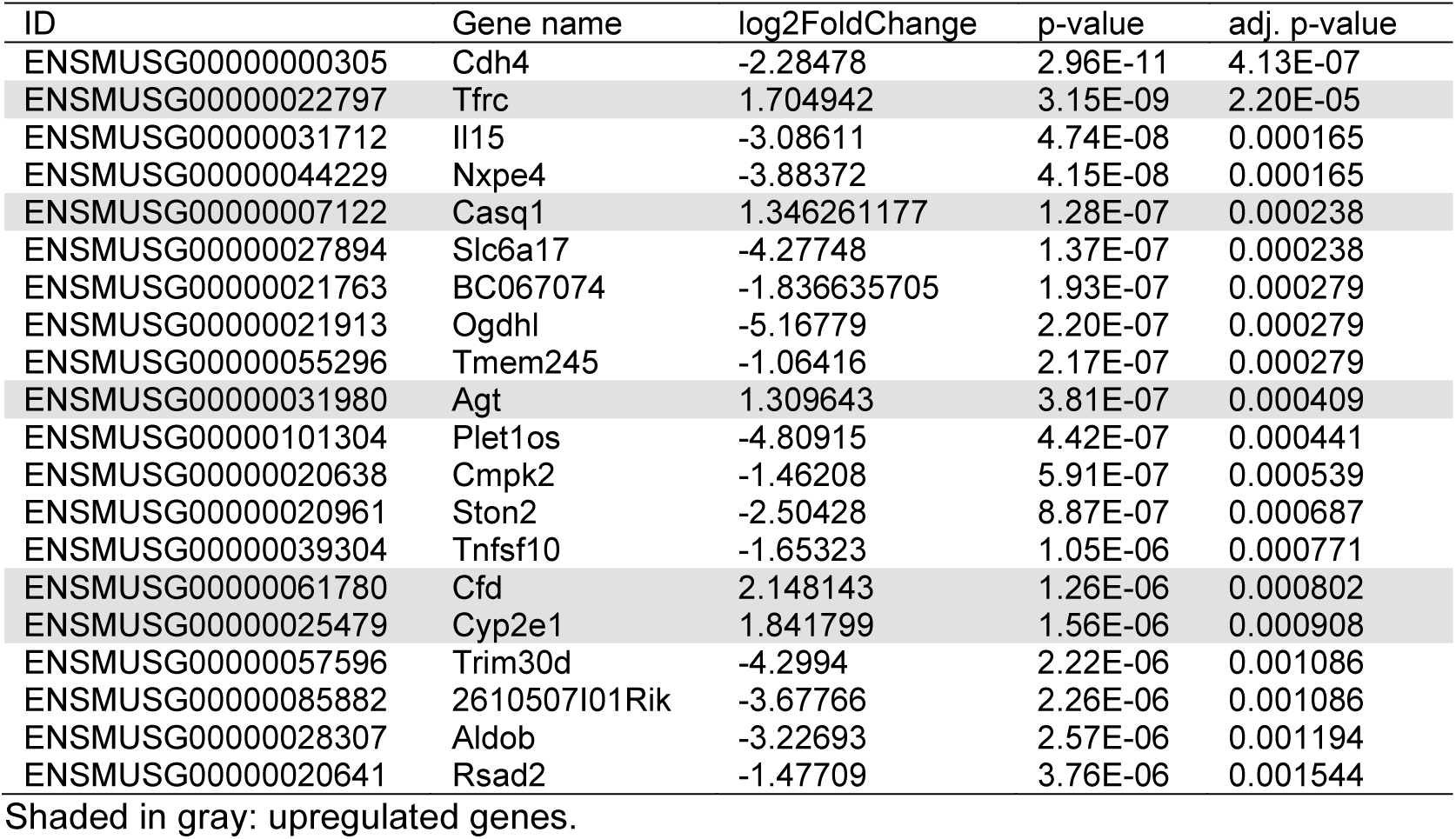
Top 20 cardiac genes significantly altered in *Il10*^-/-^ mice (*vs*. control).

### 3.5. FMT altered cardiac transcriptome profiles in both DSS and Il10^-/-^ mice

Since FMT rescued colitis and colitis-induced cardiac impairment (Figure 2), we further analyzed whether and how FMT alters the cardiac transcriptomic profiles in DSS-treated and *Il10^-/-^* mice using RNA-seq. PCA plot showed that FMT-treated DSS mice didn’t cluster well, suggesting variabilities between the samples (Figure 4A). Volcano plot (Figure 4B) and Hierarchal clustering (Figure 4C) of DEGs revealed 3 significant DEGs of which *Map3k6* and *Gck* were upregulated, and *Fos* was downregulated by FMT when compared to DSS mice. FMT-treated *Il10^-/-^* mice were better grouped in PCA plot (Figure 4D). According to the volcano plot, FMT-treated *Il10^-/-^* mice had 67 DEGs, among which 34 genes were upregulated, and 33 genes were downregulated compared to vehicle-treated *Il10^-/-^* mice (Figure 4E, Table S9). The top 20 DEGs are listed in Table 3 and the top 30 DEGs are displayed in a heat map (Figure 4F). The top 5 upregulated cardiac genes by FMT include *Prox1*, *Kctd12b*, *Dach1*, *Itgb6*, and *Nav3* and the top 5 downregulated genes are *Ciart1*, *Cdkn1a*, *Mt1*, *Mmp3*, and *Ddit4* (Table 3). KEGG pathway analysis of the 67 significant DEGs revealed 10 enriched terms with significant p-values (Table S10). Dysregulation of p53 signaling pathway, transcriptional misregulation in cancer, human papillomavirus infection, focal adhesion, PI3K-Akt signaling pathway, lipid and atherosclerosis, IL-17 signaling, and NF-kappa B signaling pathways was observed. GO enrichment analysis revealed enrichments in 260 biological processes (Table S11). Top 10 enriched GO biological processes include regulation of neuroinflammatory response, regulation of cell motility and migration, regulation of striated/non-striated cardiac muscle cell and epithelial cell apoptotic processes, and regulation of DNA and bile acid biosynthetic processes. These results suggest that FMT could rescue multiple dysregulated signaling pathways in the hearts of mice with DSS-induced chronic colitis and spontaneous colitis.

**Fig. 4.**
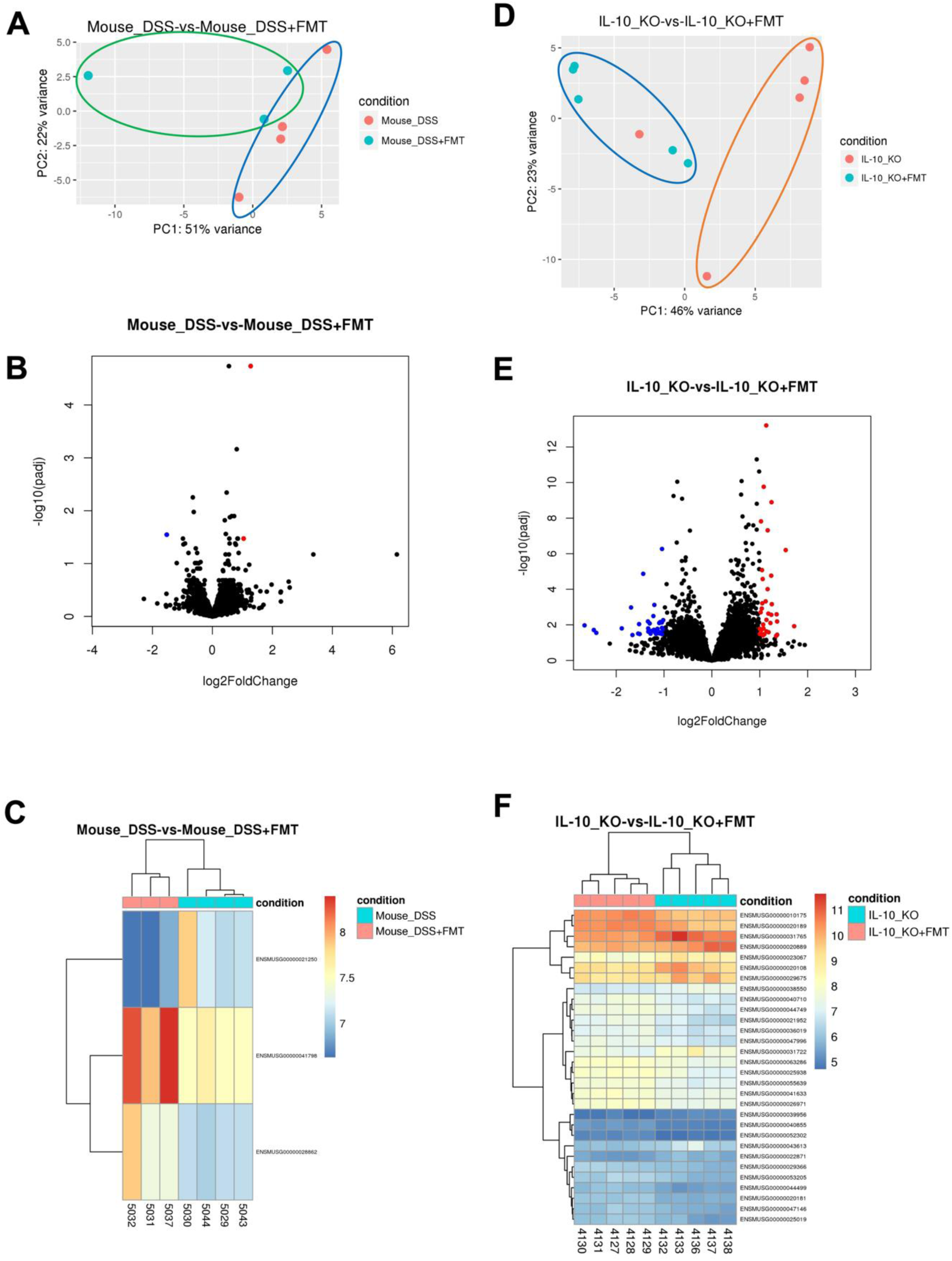
Genome-wide survey of cardiac mRNAs rescued by FMT in DSS and *Il10*^-/-^ mice. (***A***) Principal component analysis (PCA) of DSS+FMT mice (turquoise circles) *vs*. DSS mice (orange circles). (***B***) Volcano plot of differential expression analysis of cardiac mRNAs in DSS+FMT mice *vs*. DSS mice. Blue circles represent downregulated genes, and red circles represent upregulated genes. (***C***) Heat map showing top 3 cardiac genes rescued by FMT. (***D***) PCA analysis of *Il10*^-/-^ +FMT *vs*. *Il10*^-/-^ mice. (***E***) Volcano plot of differential expression analysis of cardiac mRNAs altered by FMT (*Il10*^-/-^+FMT *vs*. *Il10*^-/-^ mice). (***F***) Heat map showing top 30 cardiac genes rescued by FMT in *Il10*^-/-^ mice. n=3 or 4.

**Table 3.** Top 20 cardiac genes significantly altered by FMT in *Il10*^-/-^ mice.

| ID | Gene name | log2FoldChange | p-value | Adj. p-value |
| --- | --- | --- | --- | --- |
| ENSMUSG00000010175 | Prox1 | 1.13655 | 5.08E-18 | 6.19E-14 |
| ENSMUSG00000041633 | Kctd12b | 1.083528 | 8.56E-14 | 1.74E-10 |
| ENSMUSG00000055639 | Dach1 | 1.246298 | 1.04E-12 | 1.27E-09 |
| ENSMUSG00000026971 | Itgb6 | 1.026256 | 1.61E-11 | 1.51E-08 |
| ENSMUSG00000020181 | Nav3 | 1.166006 | 7.17E-11 | 4.85E-08 |
| ENSMUSG00000038550 | Ciart | -1.04137 | 1.02E-09 | 5.40E-07 |
| ENSMUSG00000044499 | Hs3st5 | 1.542304 | 1.24E-09 | 6.28E-07 |
| ENSMUSG00000063286 | Gm8995 | 1.04713 | 2.96E-08 | 8.38E-06 |
| ENSMUSG00000023067 | Cdkn1a | -1.43546 | 5.32E-08 | 1.35E-05 |
| ENSMUSG00000040710 | St8sia4 | 1.240315 | 7.08E-08 | 1.72E-05 |
| ENSMUSG00000036019 | Tmtc2 | 1.061487 | 1.16E-07 | 2.66E-05 |
| ENSMUSG00000040855 | Reps2 | 1.164053 | 5.13E-07 | 9.91E-05 |
| ENSMUSG00000047996 | Prrg1 | 1.124163 | 3.82E-06 | 0.00048 |
| ENSMUSG00000029366 | Dck | 1.052613 | 5.12E-06 | 0.000615 |
| ENSMUSG00000044749 | Abca6 | 1.252182 | 6.13E-06 | 0.000691 |
| ENSMUSG00000031765 | Mt1 | -1.20144 | 7.01E-06 | 0.000756 |
| ENSMUSG00000043613 | Mmp3 | -1.68805 | 1.17E-05 | 0.001056 |
| ENSMUSG00000052302 | Tbc1d30 | 1.042687 | 1.40E-05 | 0.001209 |
| ENSMUSG00000020189 | Osbpl8 | 1.019306 | 2.72E-05 | 0.001979 |
| ENSMUSG00000047146 | Tet1 | 1.179904 | 3.43E-05 | 0.002335 |

Shaded in gray: upregulated genes.

### 3.6. Quantitative analysis of dysregulated cardiac genes by RT-qPCR

To verify some DEGs identified by bulk RNA-seq in both DSS colitis and *Il10^-/-^* mice, we performed RT-qPCR with *Gapdh* as the housekeeping gene. Indeed, significantly increased mRNA expression was confirmed for *Gdpd3* (Figure 5A), *Ppm1k* (Figure 5B), *Fos* (Figure 5C), and *Ctgf* (Figure 5D) in DSS-treated mice compared to controls. Decreased mRNA expression of *Zfp970* (Figure 5E), *Serpine1* (Figure 5F), *Ano5* (Figure 5G), and *Myot* (Figure 5H) was also verified in DSS mice compared to controls. Furthermore, healthy control donor FMT ameliorated the upregulation of *Gdpd3*, *Ppm1k*, *Fos*, and *Ctgf* and the downregulation of *Zfp970*. Though, FMT showed no mitigation of decreased mRNA expression of *Serpine1*, *Ano5*, and *Myot* compared to vehicle-treated DSS mice (Figures 5A-5H). In *Il10^-/-^* mice *vs*. WT mice, mRNA overexpression of *Casq1* (Figure 5I), *Il1b* (Figure 5J), *Agt* (Figure 5K), and *Trfc* (Figure 5L) were confirmed by RT-qPCR. Additionally, reduced mRNA expression of *Nxpe4* (Figure 5M), *Slc6a17* (Figure 5N), and *Ogdhl* (Figure 5O) were detected. Furthermore, WT mice donor FMT significantly ameliorated the elevation of *Casq1*, *Il1b*, and *Agt* and slightly decreased *Trfc* overexpression, though not statistically significant. FMT was unable to mitigate the reduction of *Nxpe4*, *Slc6a*, and *Ogdhl* in *Il10^-/-^* mice (Figures 5I-5O). Gene transcription patterns observed by RNA-seq are directly supported by RT-qPCR experiments, reinforcing the presence of transcriptional alterations at the mRNA level.

**Fig. 5.**
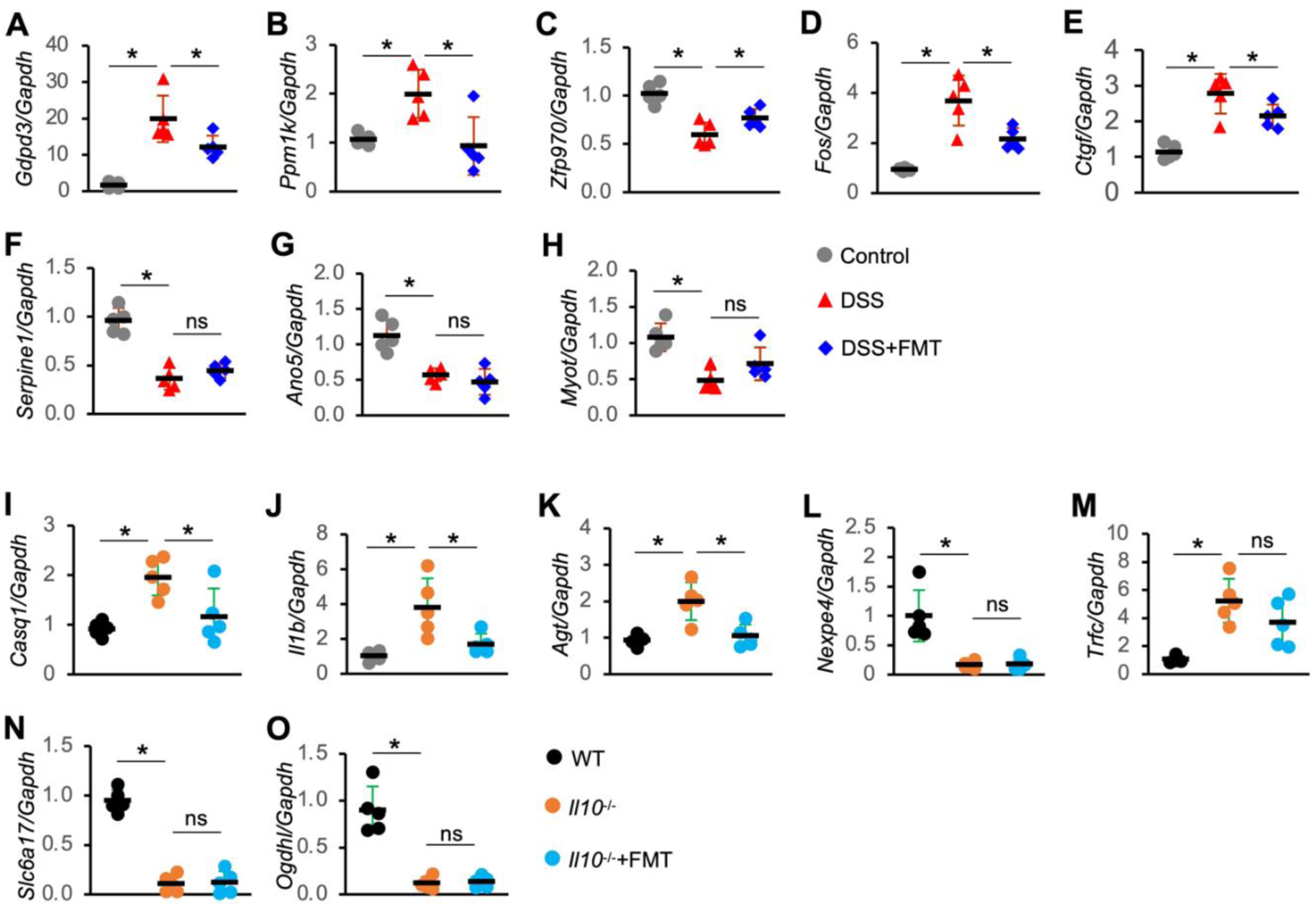
Verification of mRNA levels of selected genes in the hearts of DSS and *Il10*^-/-^ mice. mRNA expression was detected by RT-qPCR and *Gapdh* served as the internal control. n=5. One-way ANOVA. * *p*<0.05. ns, not significant.

### 3.7 Colitis-induced cardiac fibrosis was markedly mitigated by FMT

Myocardial fibrosis is a common pathophysiological companion of many cardiac disorders. To investigate if chronic colitis promotes fibrosis in the adult heart, we evaluated selected proteins that are related to fibrosis using WB. Fibrotic proteins collagen type 1 alpha 1 (COL1A1), COL3A1, and fibronectin were significantly upregulated in the hearts of DSS and *Il10^-/-^* mice (Figure 6A), indicative of enhanced cardiac fibrosis. Masson’s trichrome staining revealed more deposited collagen fibers in both mouse models (Figure 6B), confirming that chronic colitis promotes fibrosis. FMT significantly mitigated the increases of fibrotic proteins and collagen fibers in DSS mice. Similar but less effect was observed in *Il10^-/-^* mice. Profibrotic protein, β-Catenin (encoded by *CTNNB1* gene) and vascular endothelial cadherin (vE-cadherin) expression was also increased in both DSS and *Il10^-/-^*mice, but FMT only significantly ameliorated the increase in DSS-treated mice, not in *Il10^-/-^* mice (Figure 6A). Additionally, B-cell lymphoma 2 (BCL-2) and Cyclin D1 (CCND1), two known downstream targets of β-Catenin, as well as staphylococcal nuclease domain-containing 1 (SND1), were upregulated in DSS mice and the upregulation was mitigated by FMT. BCL-2 and SND1 expression showed marginal differences among WT, *Il10^-/-^*, and *Il10^-/-^*+FMT mice while Cyclin D1 showed increased expression only after FMT. Finally, DNA methyltransferase 3 alpha (DNMT3A) expression was suppressed in the hearts of DSS mice; the suppression was mitigated by FMT. DNMT3A expression remained unchanged in *Il10^-/-^* mice compared to WT mice and FMT slightly elevated DNMT3A levels.

**Fig. 6.**
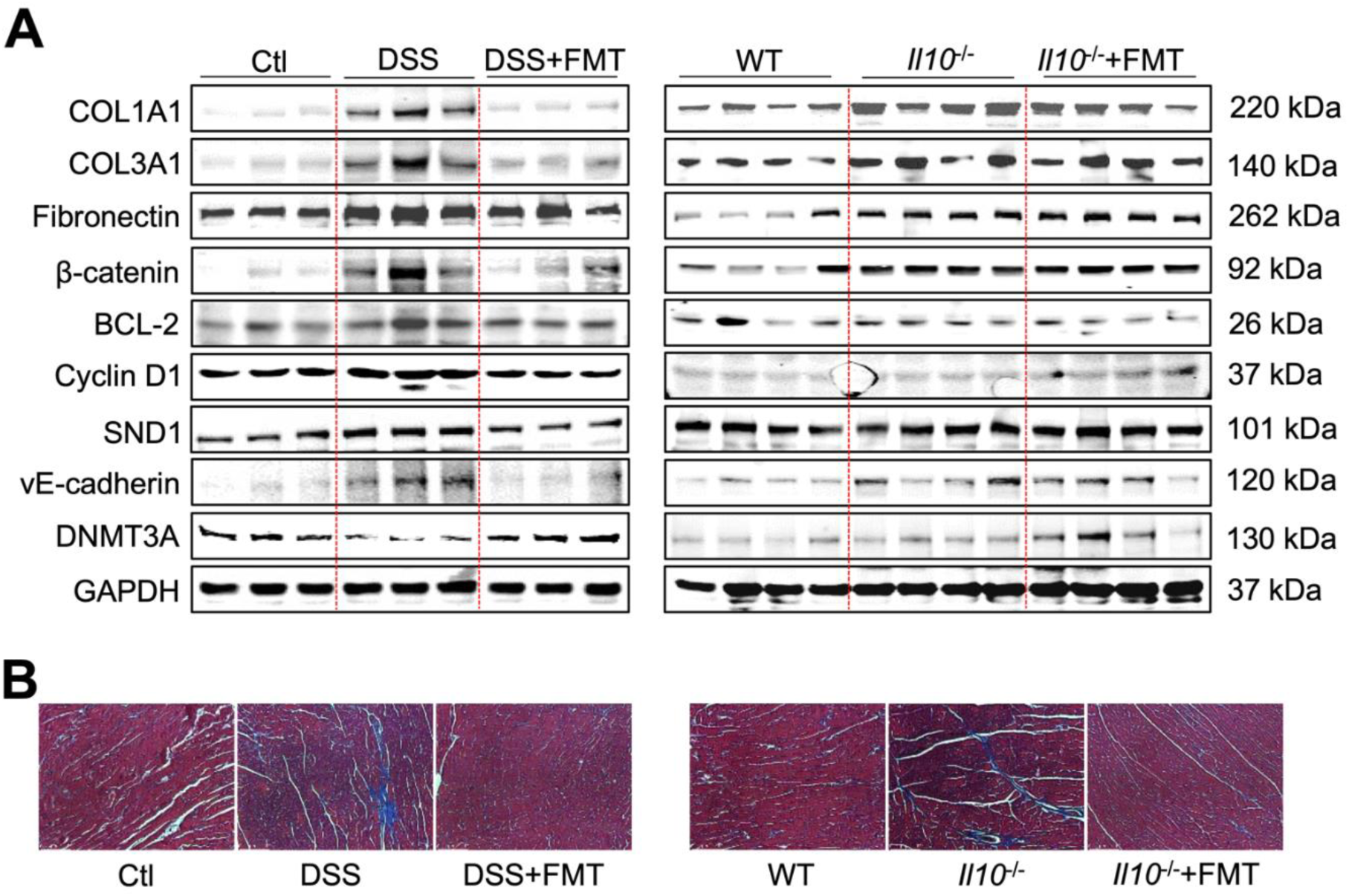
FMT mitigated multiple cardiac proteins dysregulated by chronic colitis in DSS and *Il10*^-/-^ mice. (***A***) Western blot analysis of selected proteins. Total tissue lysates were prepared from frozen heart tissue. GAPDH served as a loading control. (***B***) Masson’s trichrome staining of fixed heart tissue (10X magnification).

## 4. Discussion

Reports of an increased prevalence of cardiac complications in patients with IBD have accumulated over recent years and IBD prevalence is rising with extraintestinal manifestations presenting in 25-40% of patients. However, the molecular pathogenic mechanisms of cardiovascular derangements in IBD remain poorly understood (1, 2). Our findings demonstrate that chronic colitis induces significant functional cardiac impairment, characterized by molecular remodeling and cardiac fibrosis in DSS-treated and *Il10*^-/-^ mice. Notably, therapeutic FMT was shown to alleviate cardiac fibrosis, remodeling, and functional impairment even in the absence of anti-inflammatory protein, IL-10. Differentially expressed genes identified through bulk RNA-seq were validated by RT-qPCR, and dysregulated protein expression was confirmed using WB analysis.

Chronic colitis leads to significant cardiac dysfunction and molecular remodeling, as evidenced histologically by cardiac fibrosis and inflammation. Supportive to this phenotype, increased fibrotic protein (i.e. COL1A1, COL3A1, Fibronectin) and pro-fibrotic protein (i.e. β-catenin, vE-Cadherin) expression were observed in both DSS and *Il10*^-/-^ mice. In healthy adult cardiac tissue, constitutive expression of β-catenin remains low, lending to limited cardiomyocyte self-renewal. In response to pathogenic stimuli, like hypertrophy, Wnt/β-catenin signaling is activated, resulting in contractile dysfunction (22, 23). This phenomenon was observed in both of our models. vE-cadherin interacts with catenin proteins (like β-catenin), forming functional adhesion junctions important for modulation of endothelial cell function, vascular permeability and integrity, and lymphatic homeostasis within the heart (24, 25). Cardiac transcriptional reorganization in each model was characterized by increased transcription of fibrosis-associated genes, including cytochrome P450 2e1 (*Cyp2e1*), complement factor d (*Cfd*), mitogen-activated protein kinase kinase kinase 6 (*Map3k6*), and phosphoenolpyruvate carboxykinase 1 (*Pck1*). Cytochrome P450 enzymes metabolize fatty acids into inflammatory mediators, like eicosanoids. Prostaglandin E2, an eicosanoid, activates Wnt/β-catenin signaling, leading to E-prostanoid receptors 1-4 activation and subsequent cardiac fibroblast activation, cardiomyocyte hypertrophy/dysfunction, endothelial cell dysfunction, immune cell activation, and overall inflammation (26, 27). *Map3k6* signaling also contributes to pathological cardiomyocyte hypertrophy and fibrosis particularly observed in heart failure via extracellular signal-regulated kinase, c-Jun N-terminal kinases, and p38 signaling, and initiated by G-protein coupled receptors, receptor tyrosine kinases (e.g., insulin-like growth factor 1), receptor serine/threonine kinases (e.g., TGF-β), and cardiotrophin-1 (gp130 receptor) (28, 29). *Cfd* encodes the rate-limiting enzyme required for formation of C3 convertase in the alternative complement pathway (30). In particular, the complement C3-Cfd-C3a receptor signaling axis regulates cardiac remodeling, fibrosis, and subsequent right ventricular failure (31). The common changes observed between our two models reinforce the idea that systemic inflammation and dysbiosis induced by chronic colitis have far reaching effects on many organs across the body, importantly including the heart. However, we do not rule out additional, non-inflammatory factors, like miRNAs, as potential contributors to cardiac dysfunction in conjunction with inflammation and dysbiosis which we later discuss.

Our transcriptome analysis reveals a widely different pattern of response in the hearts of the two colitis models which we primarily attribute to lack of anti-inflammatory signaling in *Il10^-/-^* mice. In the *Il10^-/-^* model, the upregulation of *Cyp2e1* and *Pla2g4e* in the hearts draws interest due to their roles in dysregulation of long chain fatty acid metabolism – specifically arachidonic acid (AA) and linoleic acid (LA) which mediate inflammation and fibrosis. Generally, AA is a key precursor for eicosanoid synthesis, such as prostaglandins and thromboxanes (via cyclooxygenase pathways) and leukotrienes (via lipooxygenase pathways), each playing fundamental roles in regulating inflammation, immune-cell responses, and vascular changes throughout the body. In IBD, AA and LA aggravate intestinal IBD symptoms particularly due to their role in leukotriene and prostaglandin synthesis (32). Elevated dietary n-6 polyunsaturated fatty acids (n-6 PUFAs), such as AA and LA, promote myocardial fibrosis by increasing the collagen I/III ratio through eicosanoid-driven pathways and LA’s direct activation of TGF-β (33). Furthermore, explanted hearts from patients with dilated cardiomyopathy showed increased levels of metabolites derived from n-6 PUFA metabolism (34). Within each colitis model, we speculate that chronic upregulation of *Cyp2e1* transcription leads to enhanced activation and metabolism of AA and LA, which in turn promote dysregulated eicosanoid synthesis, driving cardiac inflammation and fibrosis. Furthermore, KEGG enrichment identified a dysregulation in Nod-like receptor signaling pathway genes, *Gbp3*, *Gbp4*, *Gbp5*, and *Stat2* – mediators of the inflammatory response through interferon-γ (IFN-γ) signaling mechanisms. IFN-γ promotes Th1-cell activation induced-fibrosis in nonischemic heart failure amongst other diverse inflammatory functions (35). Immunological reprogramming in the hearts of *Il10^-/-^* mice was further highlighted by KEGG and GO biological process dysregulation of the immune response, T cell proliferation, IL-18 production, granulocyte activation, and neutrophil activation, all supporting the notion that IBD-induced systemic inflammation may stimulate inflammatory responses in distant organs, including the heart.

A different gene response is revealed by molecular characterization of DSS-colitis hearts: reduced protein expression of the epigenetic regulator DNMT3A and increased expression of SND1, which encodes Tudor staphylococcal nuclease (Tudor-SN), BCL-2, and Cyclin D1 in mouse hearts. DNMT3A inactivation induces cardiac fibrosis via enhanced release of heparin-binding epidermal growth factor-like growth factor, resultantly activating cardiac fibroblasts (36). Tudor-SN signaling promotes heart regeneration after cardiac injury via Hippo signaling pathway inactivation and subsequent Yes-associated protein (YAP) nuclear translocation and transcription in neonates (37). Downstream transcriptional programing from YAP activation facilitates fibroblast activation and fibrotic protein deposition in cardiac tissue as well as many other tissues across the body (38–40). Tudor-SN also enhances angiotensin II type 1 receptor expression, commonly appreciated for its role in cardiovascular disease development (41). Cardiac transcriptomes were characterized by upregulation of fibrotic and profibrotic transcription factors such as *Col14a1*, *Col11a1*, *Col4a6*, natriuretic peptide precursor A (*Nppa*), secreted frizzled-related protein 2 (*Sfrp2*), ECM modulator A disintegrin and metalloprotease 19 (*Adam19*), and early growth response factor 1 (*Egr1*). Additionally, DSS-colitis hearts showed KEGG pathway enrichment in cAMP signaling and lipolysis in adipocytes, along with GO biological process enrichment in cGMP-mediated signaling, tau-protein kinase activity, metal/sodium ion transport, and positive regulation of metabolic processes. These findings highlight disruptions in intracellular and intercellular signaling, as well as altered energy expenditure, in response to colitis-induced systemic changes. Importantly, DSS mice harbor an intact anti-inflammatory response, which may inherently mitigate cardiac fibrosis and dysfunction compared to *Il10^-/-^*mice. This raises the question of whether non-inflammatory pathway dysregulation could also contribute to cardiac fibrosis in DSS mice. Pursuant to this idea, we observed dysregulation of Cushing syndrome (CS)-associated genes like Ca^2+^/calmodulin-dependent protein kinase II beta (*Camk2b*), which mediates NF-κB activation and promotes cardiac hypertrophy, and melanocortin 2 receptor accessory protein (*Mrap*), a crucial mediator in downstream ACTH receptor signaling activation (42–44). CS (hypercortisolism) is independently associated with increased cardiovascular morbidity and mortality via aberrant glucocorticoid and mineralocorticoid receptor activation leading to increased sensitivity to angiotensin II, ectopic epicardial adiposity, arterial fibrosis, and endothelial cell dysfunction (45). Direct stimulation of cardiac fibroblasts by glucocorticoid signaling induces cardiomyocyte hypertrophy, myocardial fibrosis, and subsequent CS-induced cardiomyopathy (46). Independent of inflammation, increased glucocorticoid sensitivity presents as an alternative, dysbiosis-driven mechanism inducing cardiac remodeling and dysfunction, which therapeutic FMT can target.

Our results demonstrate that FMT rescues functional cardiac impairment and reduces overall cardiac fibrosis in each mouse model, evidenced by decreased fibrotic protein expression and decreased collagen fiber deposition on Masson’s trichrome stain. These results suggest that FMT can: 1) mitigate systemic inflammation, and 2) resolve dysbiosis-related signaling dysregulation like increased cardiac corticosteroid sensitivity. Interestingly, FMT treatment in *Il10*^-/-^ mice yielded relatively less recovery from dysregulated fibrotic protein synthesis, especially Fibronectin, compared to DSS-treated mice. Additionally, in *Il10^-/-^* mice, FMT treatment had minimal impact on the protein expression of pro-fibrotic β-catenin and SND1, as well as the structural protein, vE-Cadherin. In contrast, DSS mice treated with FMT exhibited significant therapeutic recoveries of these proteins, indicating a differential response to FMT between the two models. Burrello et al. demonstrated that the therapeutic potential of FMT treatment for chronic colitis-induced intestinal inflammation is mediated through IL-10 signaling mechanisms, in particular (47). It has also been reported that IL-10 has a direct protective effect against heart, liver, and pulmonary fibrosis (48).

Additionally, AA and sphingolipid metabolism are directly regulated by IL-10 anti-inflammatory signaling (49, 50). Characterized by a compromised anti-inflammatory response, *Il10^-/-^* mice still exhibited partial phenotypic rescue, suggesting that FMT can selectively target dysbiosis-mediated pathways contributing to colitis-induced cardiac fibrosis and dysfunction, independent of IL-10. Furthermore, we speculate that non-inflammatory gene dysregulation in *Il10*^-/-^ mice is masked by profound immunologic reprogramming absent in DSS mice but remains therapeutically targetable by FMT. Previous research showed that FMT ameliorates dysbiosis and the dysregulated microbiota/short chain fatty acids/GPR41/GLP-1 axis, which directly participates in glucocorticoid-induced glycolipid metabolism disorder (51), independent of inflammation. Amelioration of increased corticosteroid sensitivity presents as mechanism by which FMT rescues cardiac fibrosis (52). Moreover, in *Il10^-/-^* mice, *Mrap*, crucial for proper functioning of the ACTH receptor (44), is downregulated by FMT treatment, supporting this idea. In DSS mice, FMT ameliorated the upregulation of *Gdpd3*, *Ppm1k*, *Fos*, and *Ctgf*, and the downregulation of *Zfp970*, as verified by RT-qPCR. Inactivation of *Fos* via SIRT3-mediated histone H3 deacetylation was demonstrated to prevent cardiac fibrosis and inflammation in mice (53). In *Il10*^-/-^ mice, we verified that FMT ameliorated the overexpression of *Casq1*, *Il1b*, and *Agt*. Though, FMT only marginally recovered the expression of *Serpine1*, *Ano5*, and *Myot* in DSS mice and *Nxpe4*, *Trfc*, *Slc6a17*, and *Ogdhl* in Il10^-/-^ mice. GO enrichment analysis revealed that FMT significantly influenced inflammatory signaling pathways, including the regulation of neuroinflammation, cardiac muscle and epithelial cell apoptosis, as well as DNA/bile acid biosynthetic processes, and cell migration/motility.

While mouse models of chronic colitis allowed us to characterize cardiac remodeling, we acknowledge the limitations inherent in using murine models. It is important to note that the genetic background of DSS-treated mice is C57BL/6J, a typically healthy strain. In contrast, the genetic background of our *Il10*^-/-^ mice is 129/Sv, which exhibits elevated baseline systemic inflammation compared to C57BL/6J mice, potentially influencing the experimental outcomes (54). We also acknowledge a relatively small sample size in our RNA-seq analysis. A larger sample size would provide greater statistical power for a more detailed examination and validation of the affected genes and pathways.

From a preventative medicine perspective, our results suggest that thorough evaluation of patients’ cardiovascular health should be conducted to establish baseline cardiac function, followed by regular monitoring. Additionally, the management of treatments and medications for IBD patients with cardiovascular disease should also be closely supervised, as certain treatments, like mesalamine and 5-ASA, have been reported to cause cardiac complications (55–57). Children with early-onset IBD have also been reported to experience cardiac dysfunction (58, 59). This underscores an important point: regardless of age, IBD can have significant effects on cardiac function. Therefore, the cardiovascular health of individuals across all age groups should be thoroughly evaluated and prioritized.

## Non-standard Abbreviations and Acronyms

5-ASA: 5-aminosalicylic acid
AA: arachidonic acid
BCL-2: B-cell lymphoma 2
BDNF: brain-derived neurotrophic factor
COL1A1: collagen type I alpha 1
COL3A1: collagen type III alpha 1
DEG: differentially expressed gene
DNMT3A: DNA methyltransferase 3 alpha
DSS: dextran sodium sulfate
FMT: fecal microbiota transplantation
FS: fractional shortening
GADPH: Glyceraldehyde 3-phosphate dehydrogenase
GO: gene ontology
IBD: inflammatory bowel disease
IFN-γ: interferon gamma
IL-10: interleukin 10
IL-1β: interleukin-1 beta
KEGG: Kyoto Encyclopedia of Genes and Genomes
LVEF: left ventricular ejection fraction
MI: myocardial infarction
MPO: myeloperoxidase
n-6 PUFA: n-6 polyunsaturated fatty acid
PCA: principle component analysis
SND1: staphylococcal nuclease domain-containing 1
TGF-β: transforming growth factor beta
TTE: transthoracic echocardiogram
Tudor-SN: Tudor-staphylococcal nuclease
WB: western blot
WT: wild type

## Acknowledgments

XSZ performed experiments and data analysis, and provided technical support; KML performed investigation and data analysis, and drafted the manuscript; SSK, ML, YX, RO, HIA, and TK performed experiments and data analysis; DWP helped with data interpretation and provided critical revision; KF helped with study concept, experimental design, and revision; QL obtained funding, conceived the project, designed the experiments, analyzed the data, prepared the figures and tables, and critically revised the manuscript..

## Supplementary Material

Tables S1-S11.

### Supplementary Data

**Table S1.**
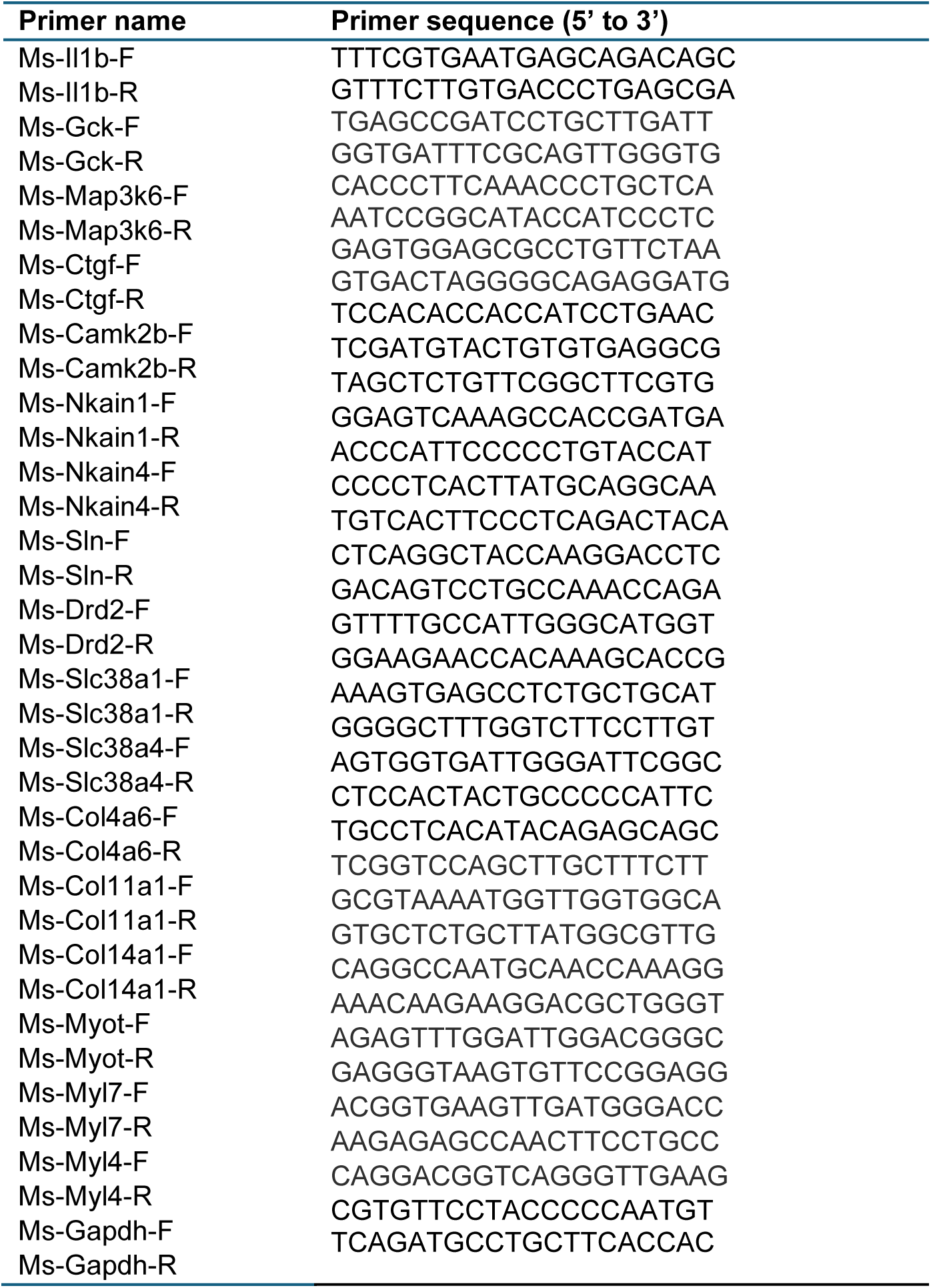
Primers for SYBR Green-based RT-qPCR.

**Table S2.** Primary antibodies used for Western blot.

| **Name of antibody** | **Catalog #** | **Manufacturer** |
| --- | --- | --- |
| anti-COL1A1 | PA5-29569 | ThermoFisher Scientific |
| anti-BCL-2 | 3498S | Cell Signaling |
| anti-GAPDH | 5174S | Cell Signaling |
| anti-COL3A1 | sc-514601 | Santa Cruz Biotechnology |
| anti-Galectin-3 | sc-32790 | Santa Cruz Biotechnology |
| anti-TIMP-1 | sc-5538 | Santa Cruz Biotechnology |
| anti-Cyclin D1 | sc-20044 | Santa Cruz Biotechnology |
| anti-vE-Cadherin | sc-52751 | Santa Cruz Biotechnology |
| anti-DNMT3A | sc-20703 | Santa Cruz Biotechnology |
| anti-β-Catenin | C2206 | Sigma-Aldrich |
| anti-SND1 | 10760-1-AP | Proteintech |
| anti-Fibronectin | ab2413 | abcam |

**Table S3.**
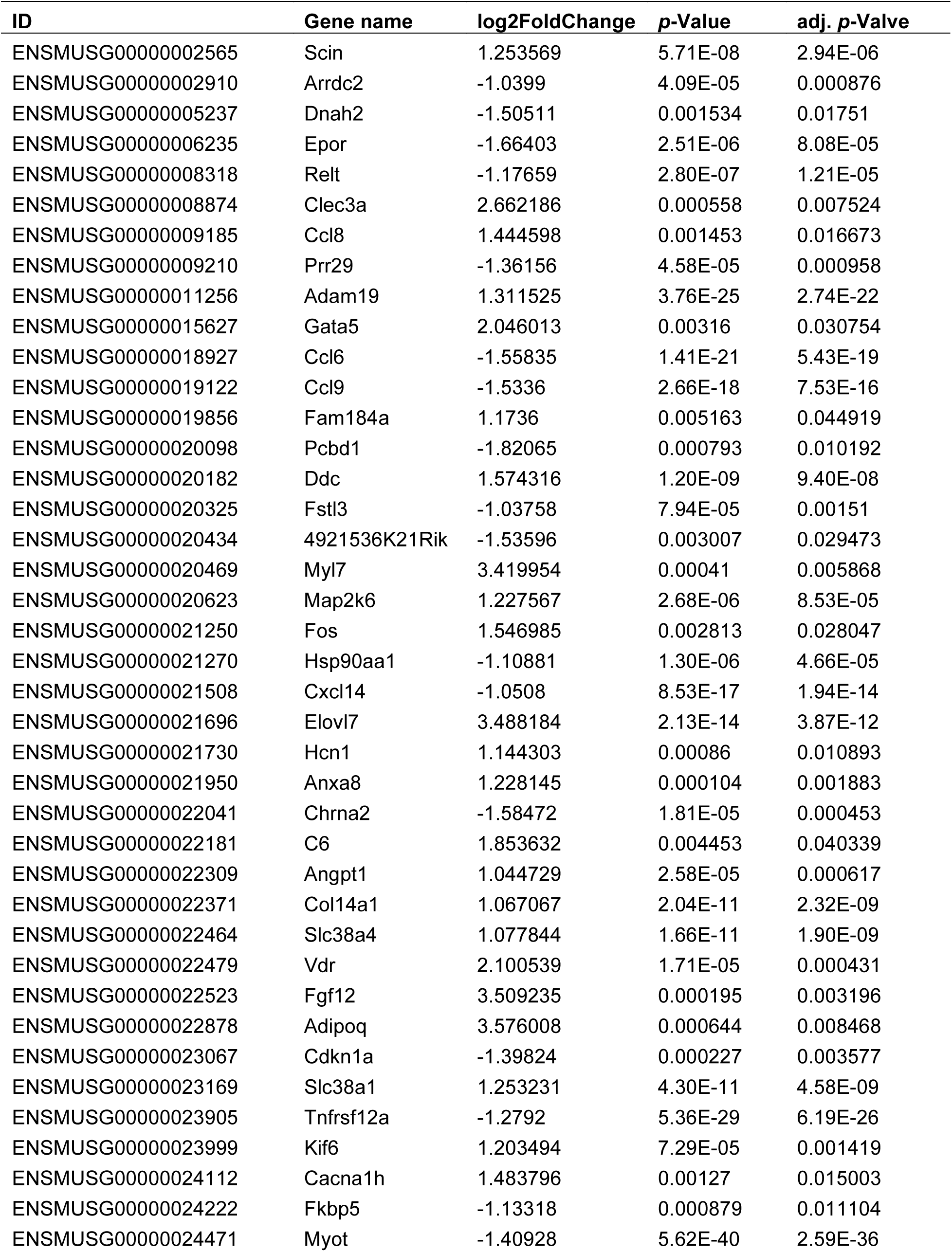
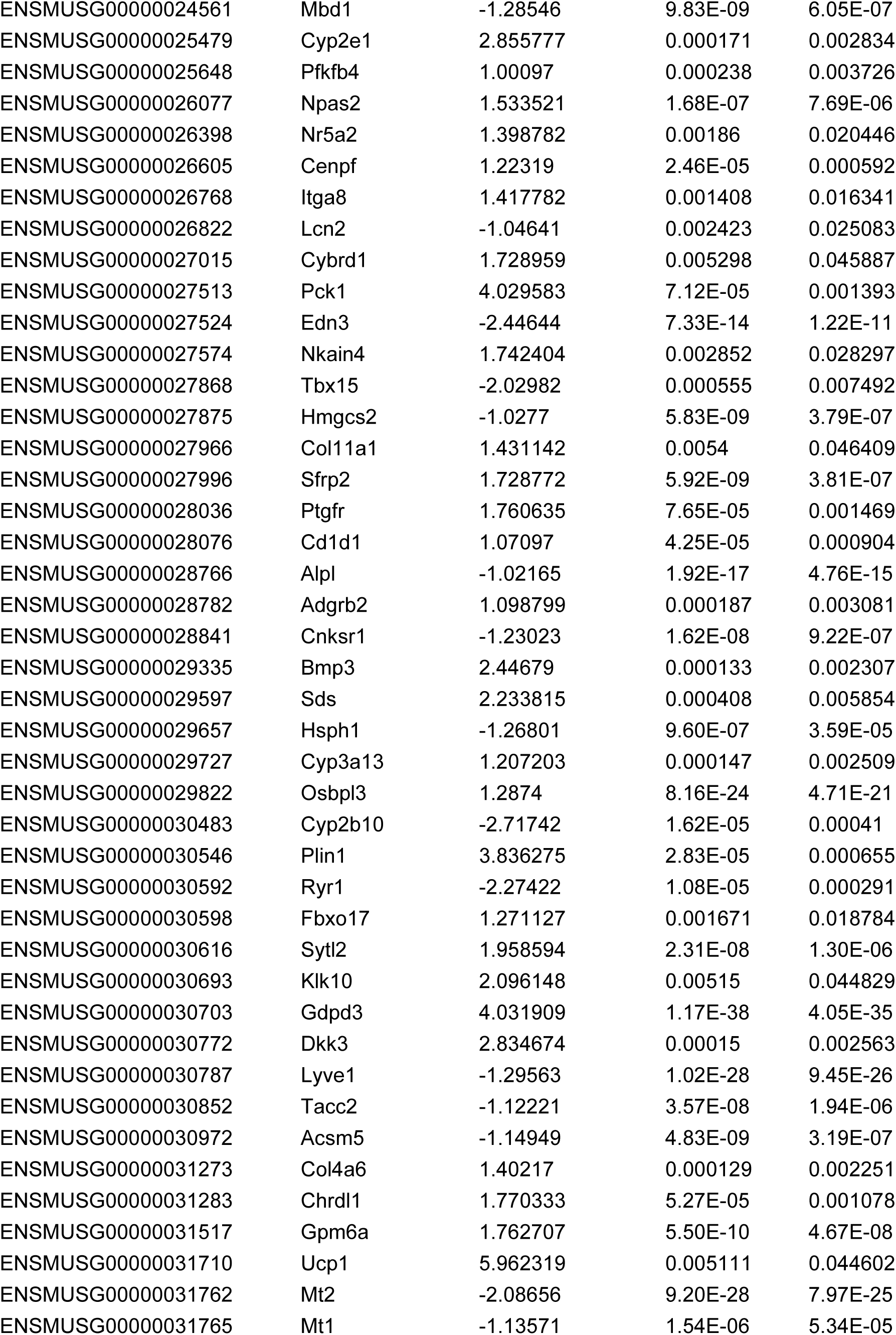
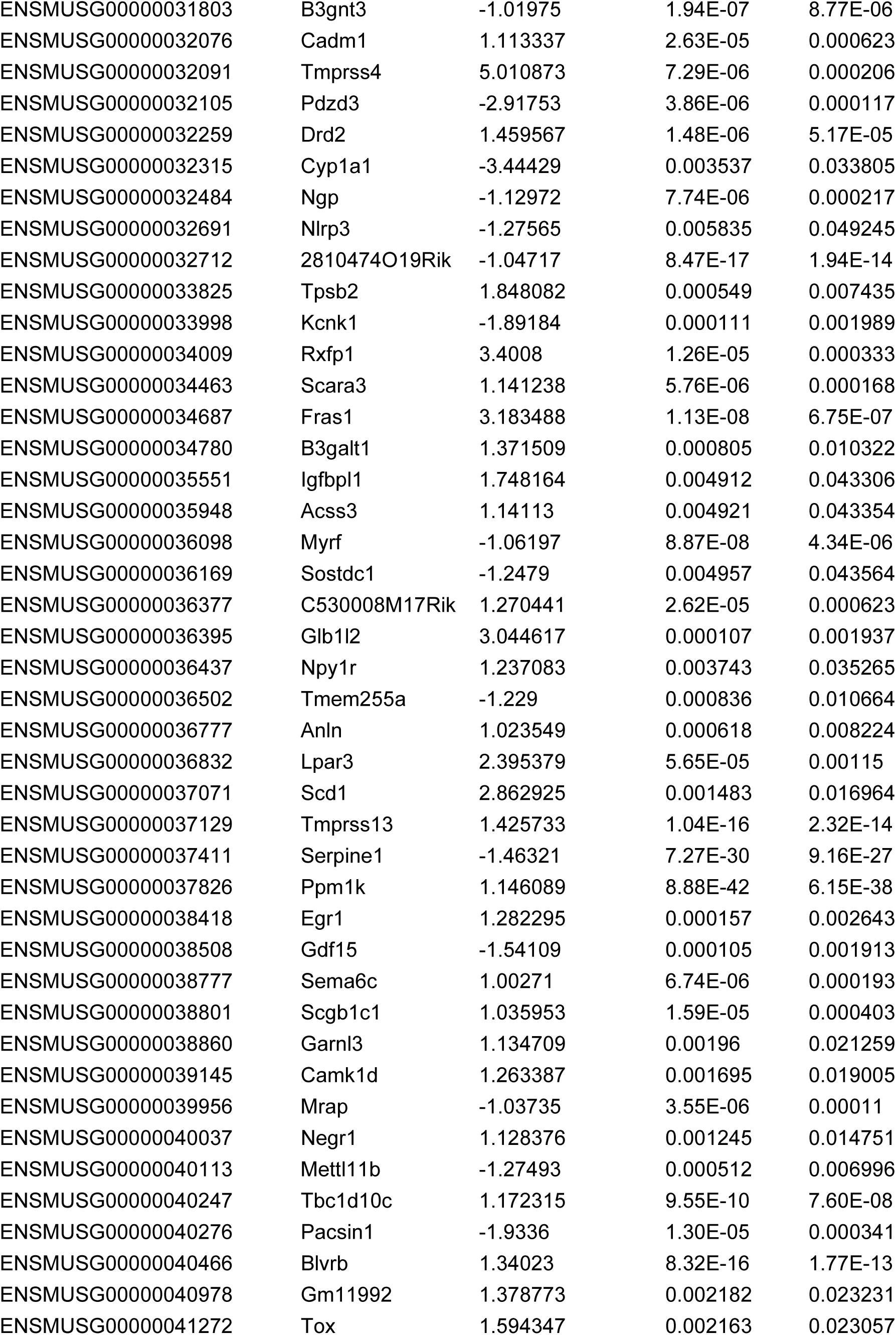
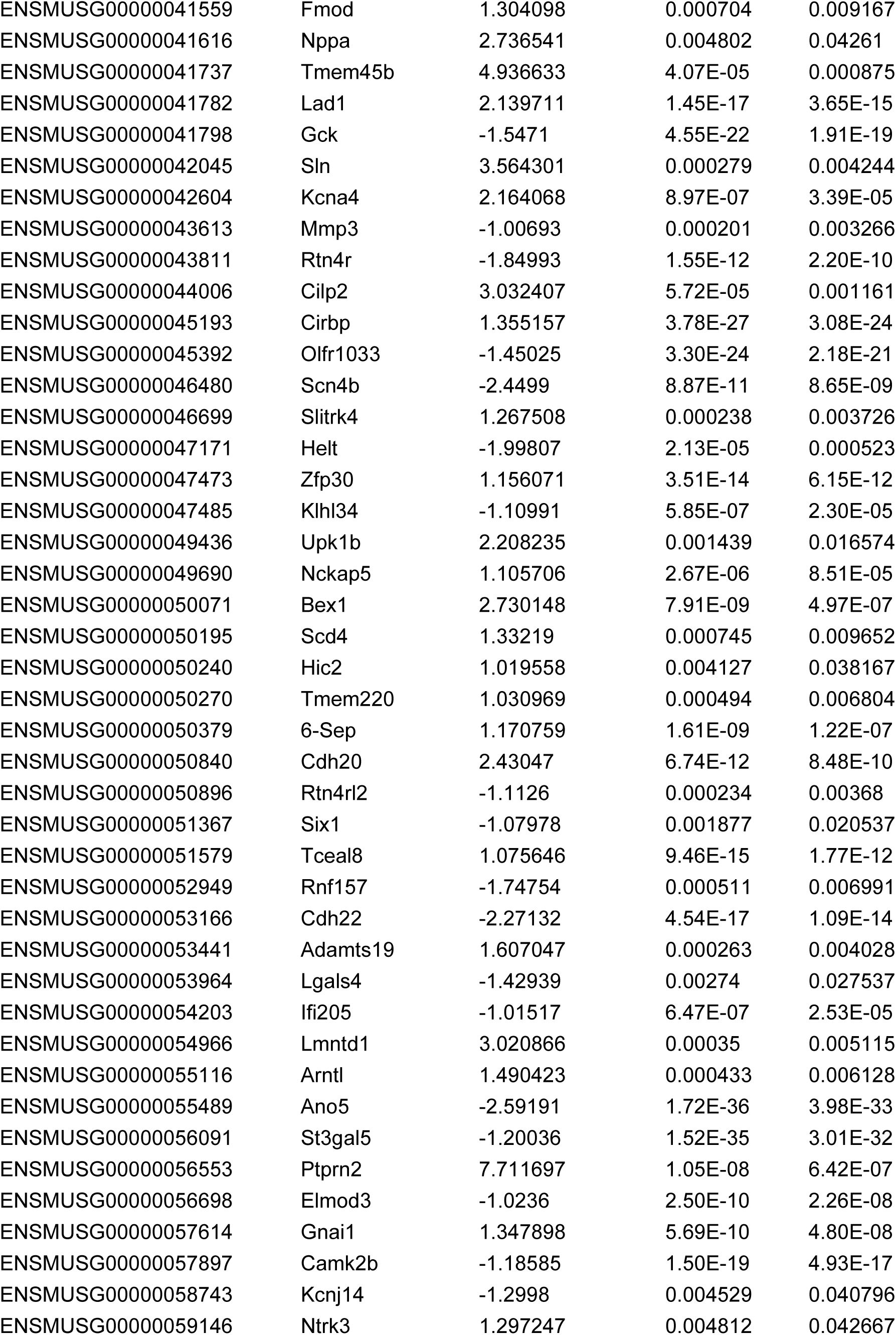
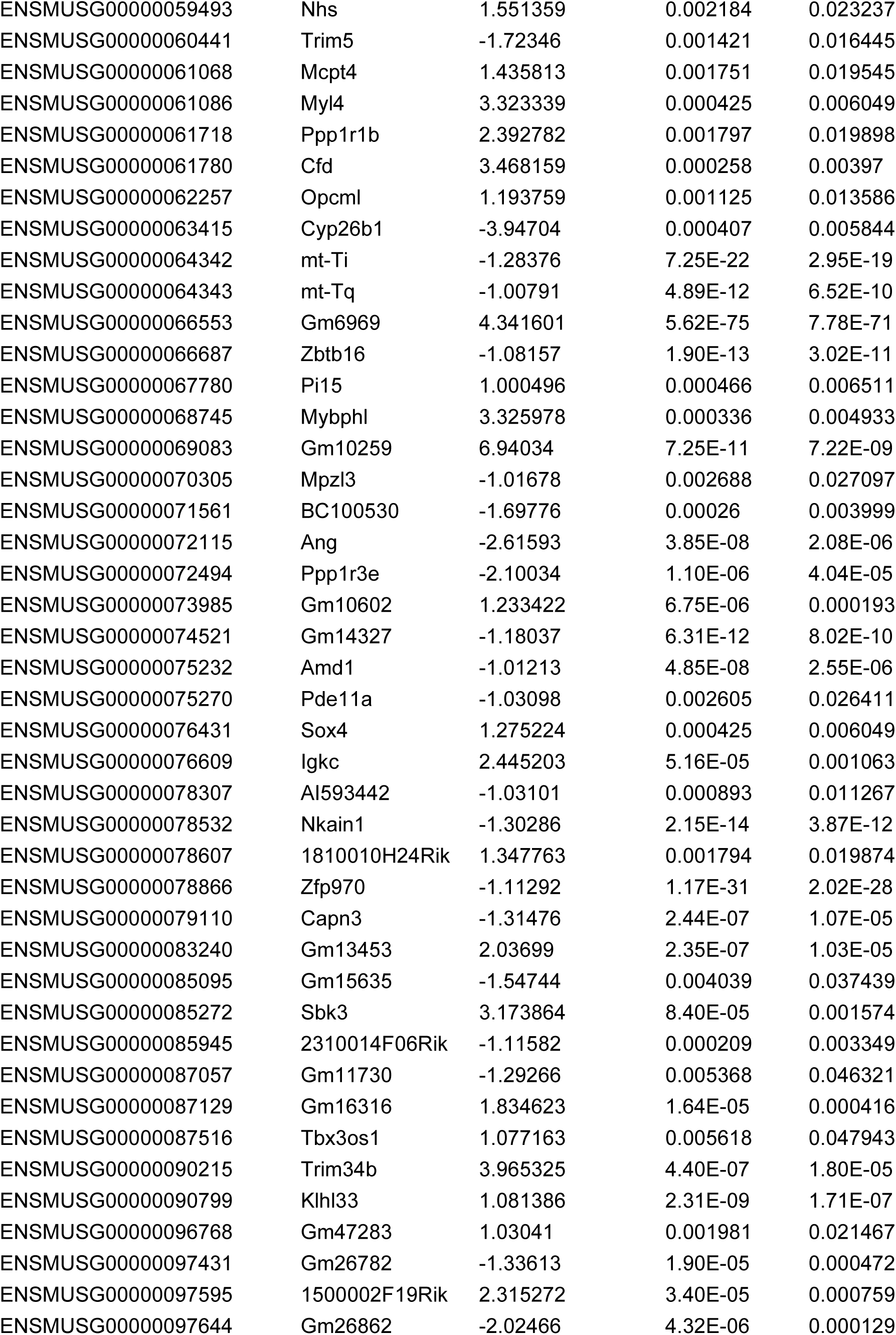
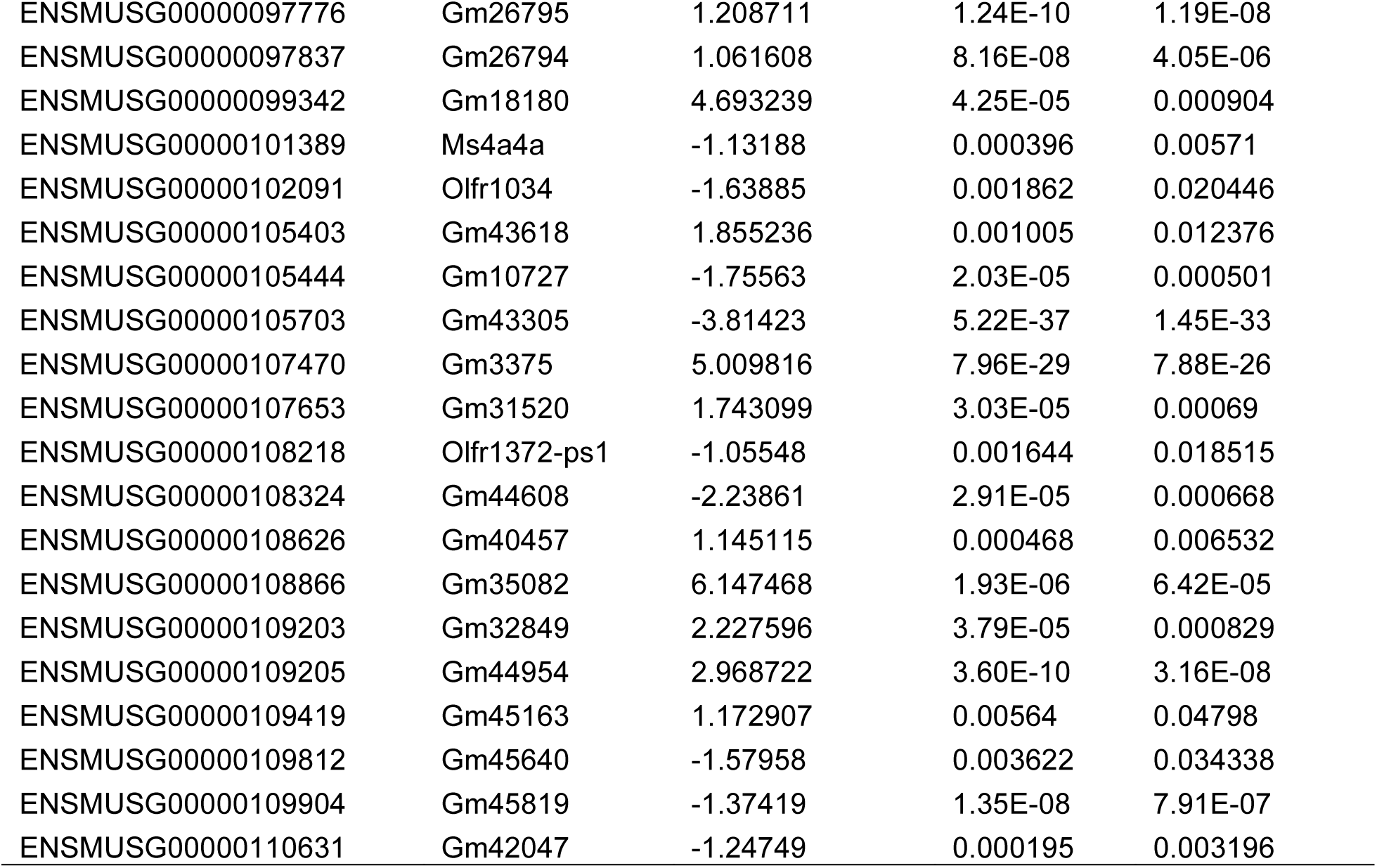
Cardiac genes significantly altered by DSS.

**Table S4.**
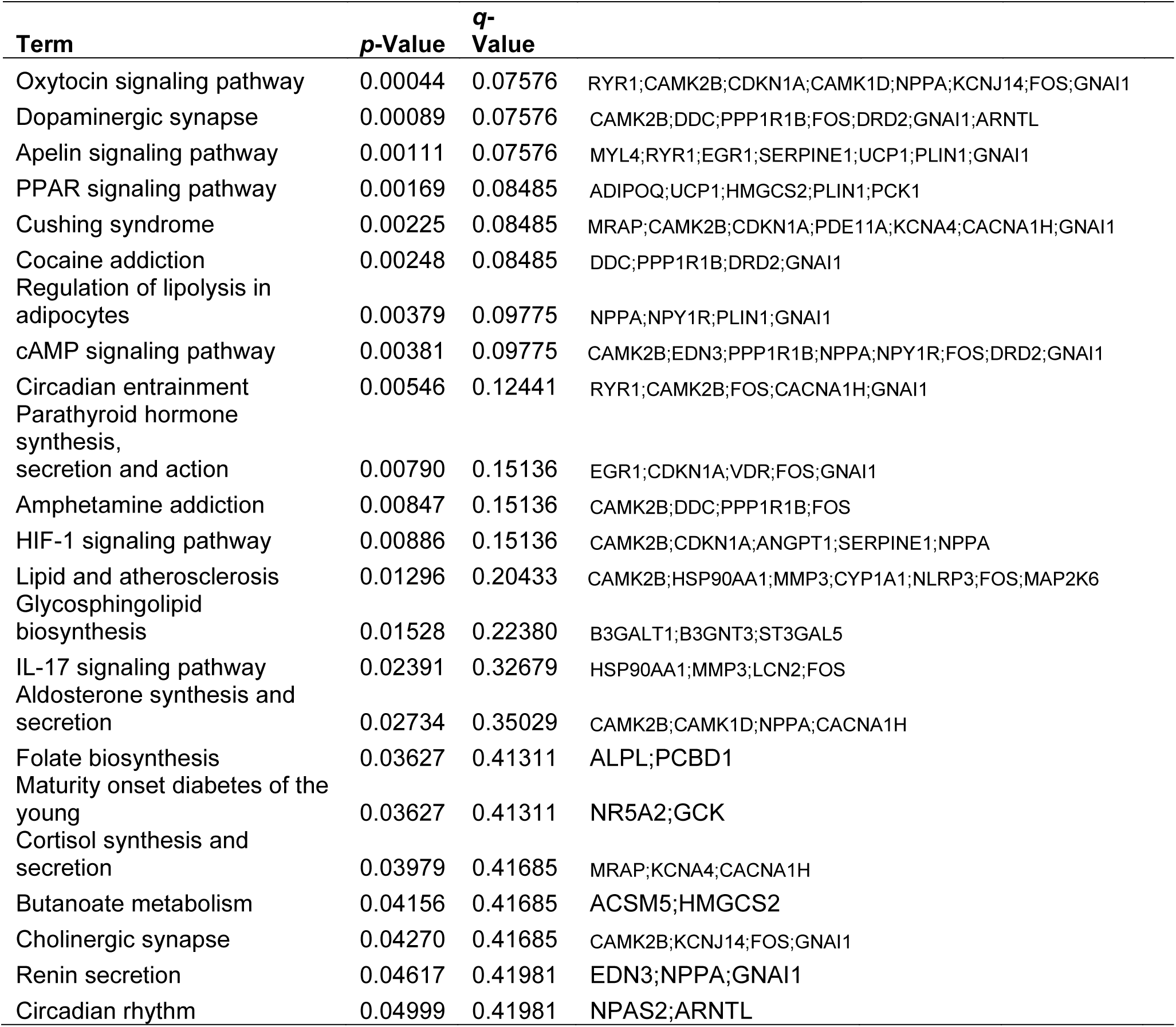
All significantly enriched signaling pathways identified by KEGG enrichment analysis (all DEGs, DSS *vs*. control).

**Table S5.** All significantly enriched biological processes identified by GO enrichment analysis (all DEGs, DSS vs. control).

| Term | p-Value | q-Value |
| --- | --- | --- |
| Regulation Of Metal Ion Transport (GO:0010959) | 4.25E-05 | 0.032568 |
| Regulation Of Sodium Ion Transport (GO:0002028) | 4.92E-05 | 0.032568 |
| Negative Regulation Of Synaptic Transmission (GO:0050805) | 0.000632 | 0.258231 |
| Positive Regulation Of Tau-Protein Kinase Activity (GO:1902949) | 0.001309 | 0.258231 |
| Ketone Body Biosynthetic Process (GO:0046951) | 0.001309 | 0.258231 |
| Regulation Of cGMP-mediated Signaling (GO:0010752) | 0.001309 | 0.258231 |
| Kidney Development (GO:0001822) | 0.001406 | 0.258231 |
| Positive Regulation Of Metabolic Process (GO:0009893) | 0.001634 | 0.258231 |
| Ketone Body Metabolic Process (GO:1902224) | 0.001949 | 0.258231 |
| Response To Alkaloid (GO:0043279) | 0.001949 | 0.258231 |
| Brown Fat Cell Differentiation (GO:0050873) | 0.003583 | 0.367371 |
| Positive Regulation Of Neuron Projection Development (GO:0010976) | 0.004366 | 0.367371 |
| Regulation Of Tau-Protein Kinase Activity (GO:1902947) | 0.004571 | 0.367371 |
| Receptor Guanylyl Cyclase Signaling Pathway (GO:0007168) | 0.004571 | 0.367371 |
| Negative Regulation Of Developmental Growth (GO:0048640) | 0.00545 | 0.367371 |
| Regulation Of Biosynthetic Process (GO:0009889) | 0.00567 | 0.367371 |
| Regulation Of Long-Term Neuronal Synaptic Plasticity (GO:0048169) | 0.00567 | 0.367371 |
| Columnar/Cuboidal Epithelial Cell Differentiation (GO:0002065) | 0.00567 | 0.367371 |
| Sarcoplasmic Reticulum Calcium Ion Transport (GO:0070296) | 0.00567 | 0.367371 |
| Positive Regulation Of Cold-Induced Thermogenesis (GO:0120162) | 0.005703 | 0.367371 |
| Hemopoiesis (GO:0030097) | 0.006469 | 0.367371 |
| Fatty Acid Biosynthetic Process (GO:0006633) | 0.006873 | 0.367371 |
| Phasic Smooth Muscle Contraction (GO:0014821) | 0.006878 | 0.367371 |
| Skeletal Muscle Fiber Development (GO:0048741) | 0.006878 | 0.367371 |
| Cellular Response To Glucose Stimulus (GO:0071333) | 0.007073 | 0.367371 |
| Regulation Of Cold-Induced Thermogenesis (GO:0120161) | 0.007209 | 0.367371 |
| Extracellular Structure Organization (GO:0043062) | 0.00886 | 0.384426 |
| External Encapsulating Structure Organization (GO:0045229) | 0.009196 | 0.384426 |
| Catecholamine Metabolic Process (GO:0006584) | 0.009606 | 0.384426 |
| Positive Regulation Of Glycogen Biosynthetic Process (GO:0045725) | 0.009606 | 0.384426 |
| Negative Regulation Of Myoblast Differentiation (GO:0045662) | 0.009606 | 0.384426 |
| Positive Regulation Of Glycogen Metabolic Process (GO:0070875) | 0.011122 | 0.384426 |
| Striated Muscle Tissue Development (GO:0014706) | 0.011122 | 0.384426 |
| Myotube Cell Development (GO:0014904) | 0.011122 | 0.384426 |
| Negative Regulation Of Blood Pressure (GO:0045776) | 0.011122 | 0.384426 |
| Heart Contraction (GO:0060047) | 0.011876 | 0.384426 |
| Regulation Of Systemic Arterial Blood Pressure (GO:0003073) | 0.012736 | 0.384426 |
| Negative Regulation Of Signal Transduction (GO:0009968) | 0.012972 | 0.384426 |
| Negative Regulation Of BMP Signaling Pathway (GO:0030514) | 0.013518 | 0.384426 |
| Intracellular Glucose Homeostasis (GO:0001678) | 0.013518 | 0.384426 |
| Positive Regulation Of Macromolecule Biosynthetic Process (GO:0010557) | 0.013943 | 0.384426 |
| Cellular Response To Ketone (GO:1901655) | 0.014386 | 0.384426 |
| Regulation Of Secretion (GO:0051046) | 0.014445 | 0.384426 |
| Regulation Of Synaptic Transmission, GABAergic (GO:0032228) | 0.014445 | 0.384426 |
| Heart Development (GO:0007507) | 0.014922 | 0.384426 |
| Metal Ion Transport (GO:0030001) | 0.014922 | 0.384426 |
| Positive Regulation Of Multicellular Organismal Process (GO:0051240) | 0.015275 | 0.384426 |
| Regulation Of Neuron Projection Development (GO:0010975) | 0.016127 | 0.384426 |
| Positive Regulation Of Cellular Biosynthetic Process (GO:0031328) | 0.016127 | 0.384426 |
| Response To Glucose (GO:0009749) | 0.016213 | 0.384426 |
| Positive Regulation Of Peptide Hormone Secretion (GO:0090277) | 0.016213 | 0.384426 |
| Regulation Of Cartilage Development (GO:0061035) | 0.016247 | 0.384426 |
| Mesenchyme Development (GO:0060485) | 0.016247 | 0.384426 |
| Regulation Of Leukocyte Chemotaxis (GO:0002688) | 0.016247 | 0.384426 |
| Positive Regulation Of Heart Rate (GO:0010460) | 0.016247 | 0.384426 |
| Triglyceride Biosynthetic Process (GO:0019432) | 0.016247 | 0.384426 |
| Cardiac Muscle Cell Contraction (GO:0086003) | 0.01814 | 0.407386 |
| Regulation Of cAMP-mediated Signaling (GO:0043949) | 0.01814 | 0.407386 |
| Regulation Of Cell Cycle G2/M Phase Transition (GO:1902749) | 0.01814 | 0.407386 |
| Glucose Homeostasis (GO:0042593) | 0.01857 | 0.410097 |
| Positive Regulation Of Fat Cell Differentiation (GO:0045600) | 0.020242 | 0.416251 |
| Pyruvate Metabolic Process (GO:0006090) | 0.020242 | 0.416251 |
| Cellular Response To Oxygen-Containing Compound (GO:1901701) | 0.020567 | 0.416251 |
| Regulation Of Protein Kinase B Signaling (GO:0051896) | 0.021288 | 0.416251 |
| SMAD Protein Signal Transduction (GO:0060395) | 0.021327 | 0.416251 |
| Adenylate Cyclase-Inhibiting G Protein-Coupled Receptor Signaling Pathway (GO:0007193) | 0.022444 | 0.416251 |
| Cardiac Muscle Tissue Development (GO:0048738) | 0.022444 | 0.416251 |
| Positive Regulation Of Heart Contraction (GO:0045823) | 0.024339 | 0.416251 |
| Acylglycerol Biosynthetic Process (GO:0046463) | 0.026571 | 0.416251 |
| Regulation Of Cardiac Muscle Cell Membrane Repolarization (GO:0099623) | 0.026571 | 0.416251 |
| Negative Regulation Of Response To External Stimulus (GO:0032102) | 0.027841 | 0.416251 |
| Striated Muscle Contraction (GO:0006941) | 0.028496 | 0.416251 |
| Response To Organic Cyclic Compound (GO:0014070) | 0.029801 | 0.416251 |
| Renal System Development (GO:0072001) | 0.029801 | 0.416251 |
| Organic Substance Transport (GO:0071702) | 0.030102 | 0.416251 |
| Regulation Of Chondrocyte Differentiation (GO:0032330) | 0.033736 | 0.416251 |
| Regulation Of Glycogen Biosynthetic Process (GO:0005979) | 0.033736 | 0.416251 |
| Activation Of Protein Kinase B Activity (GO:0032148) | 0.033736 | 0.416251 |
| Negative Regulation Of Response To Wounding (GO:1903035) | 0.033736 | 0.416251 |
| Regulation Of Synapse Organization (GO:0050807) | 0.033736 | 0.416251 |
| Cellular Response To Hexose Stimulus (GO:0071331) | 0.033736 | 0.416251 |
| Long-Chain Fatty Acid Biosynthetic Process (GO:0042759) | 0.033736 | 0.416251 |
| Cellular Response To Salt (GO:1902075) | 0.038215 | 0.416251 |
| Monocarboxylic Acid Biosynthetic Process (GO:0072330) | 0.039794 | 0.416251 |
| Calcium Ion Transport (GO:0006816) | 0.041554 | 0.416251 |
| Regulation Of Sodium Ion Transmembrane Transport (GO:1902305) | 0.041559 | 0.416251 |
| Cellular Response To Osmotic Stress (GO:0071470) | 0.041559 | 0.416251 |
| Fatty Acid Derivative Biosynthetic Process (GO:1901570) | 0.041559 | 0.416251 |
| Cardiac Muscle Cell Action Potential (GO:0086001) | 0.044304 | 0.416251 |
| Regulation Of Protein Modification By Small Protein Conjugation Or Removal (GO:1903320) | 0.044304 | 0.416251 |
| Positive Regulation Of Neural Precursor Cell Proliferation (GO:2000179) | 0.044304 | 0.416251 |
| Regulation Of Monoatomic Ion Transmembrane Transporter Activity (GO:0032412) | 0.046173 | 0.416251 |
| Negative Regulation Of Carbohydrate Metabolic Process (GO:0045912) | 0.047114 | 0.416251 |
| Negative Regulation Of Axon Extension (GO:0030517) | 0.047114 | 0.416251 |
| Cardiac Muscle Cell Action Potential Involved In Contraction (GO:0086002) | 0.047114 | 0.416251 |
| Positive Regulation Of Protein Modification Process (GO:0031401) | 0.047747 | 0.416251 |
| Positive Regulation Of Cell Projection Organization (GO:0031346) | 0.048711 | 0.416251 |
| Negative Regulation Of Cellular Response To Growth Factor Stimulus (GO:0090288) | 0.04954 | 0.416251 |
| acyl-CoA Metabolic Process (GO:0006637) | 0.049987 | 0.416251 |
| Regulation Of Transmembrane Receptor Protein Serine/Threonine Kinase Signaling Pathway (GO:0090092) | 0.049987 | 0.416251 |
| Positive Regulation Of Leukocyte Migration (GO:0002687) | 0.049987 | 0.416251 |
| Negative Regulation Of DNA Biosynthetic Process (GO:2000279) | 0.049987 | 0.416251 |

**Table S6.**
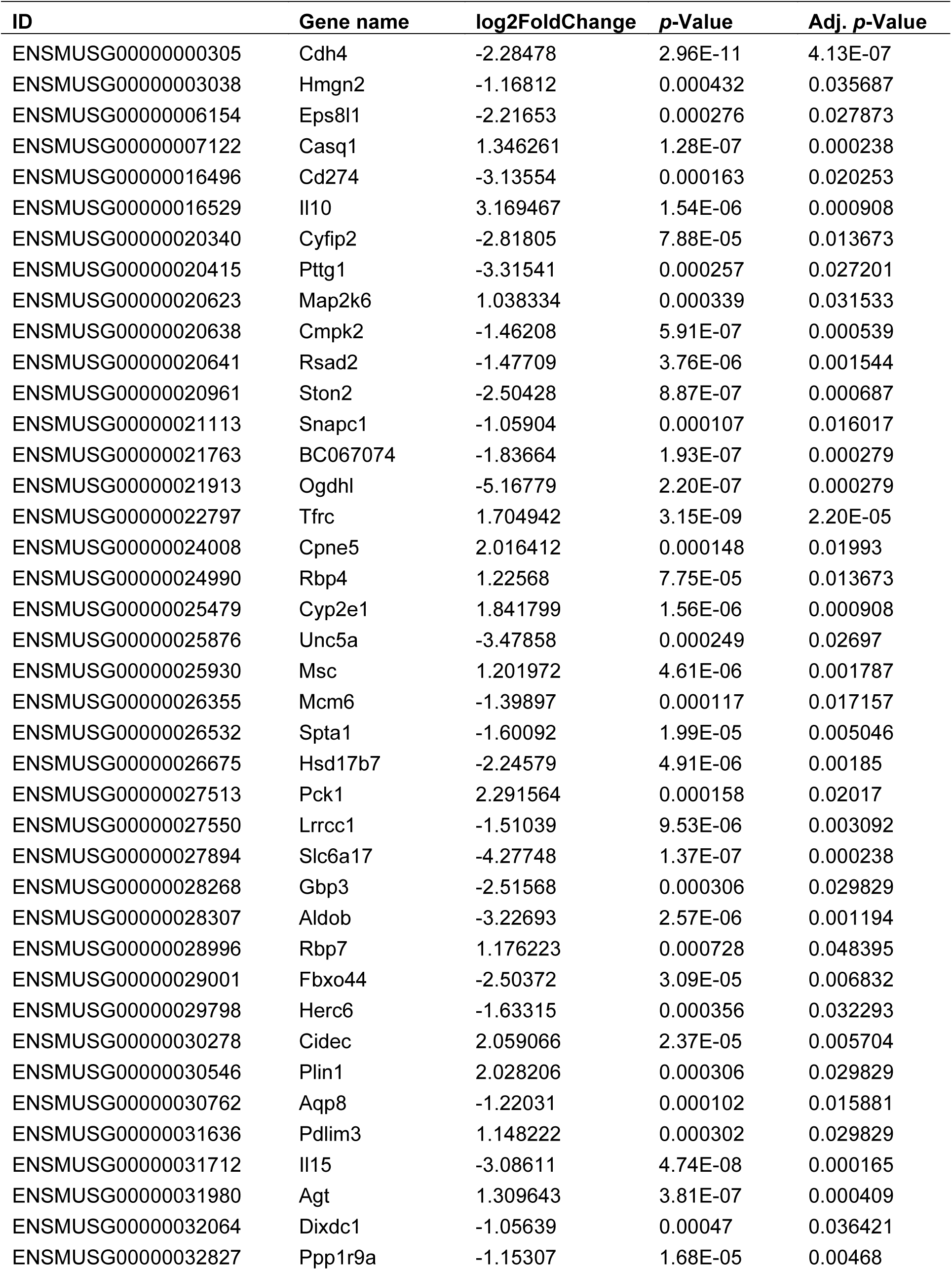
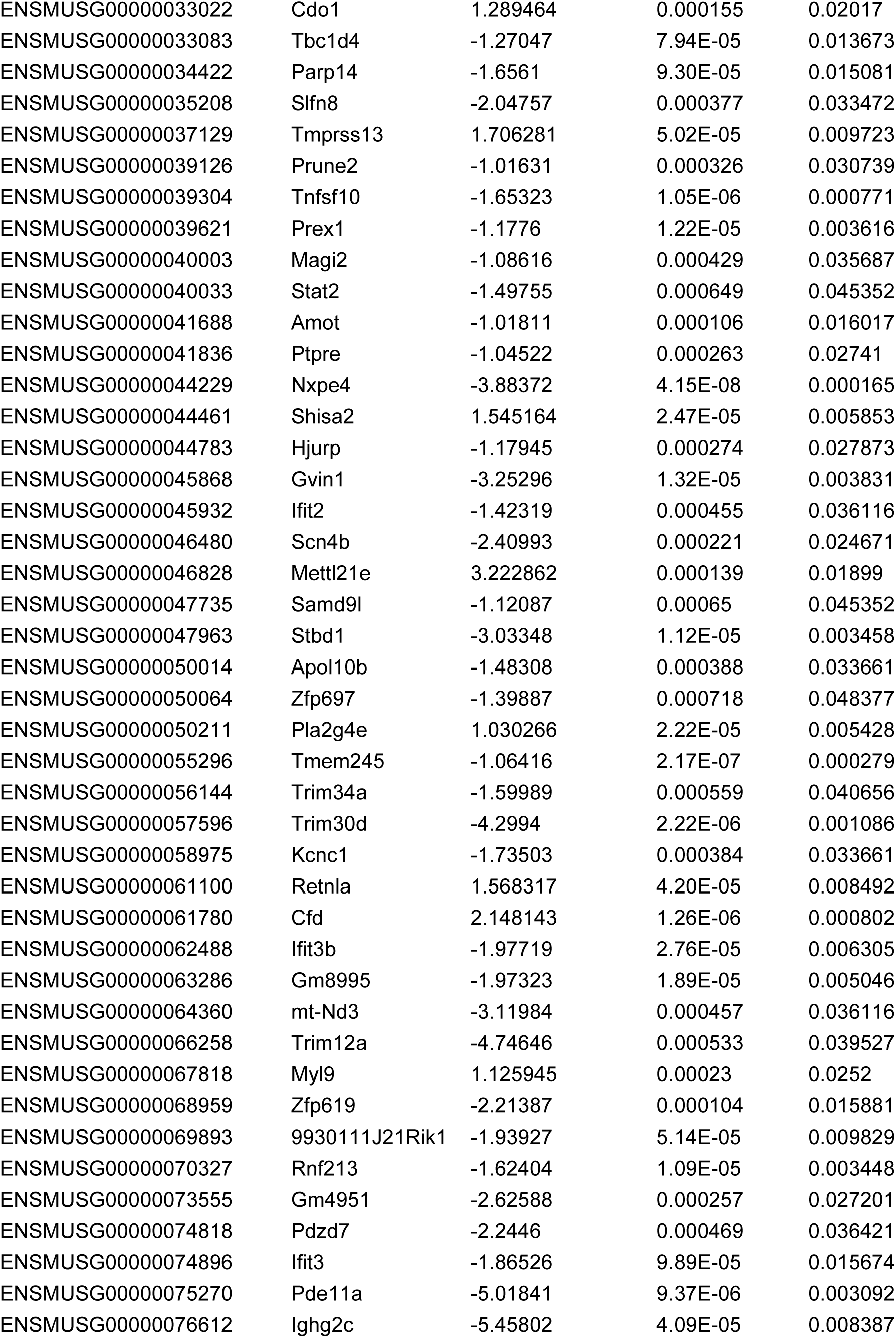
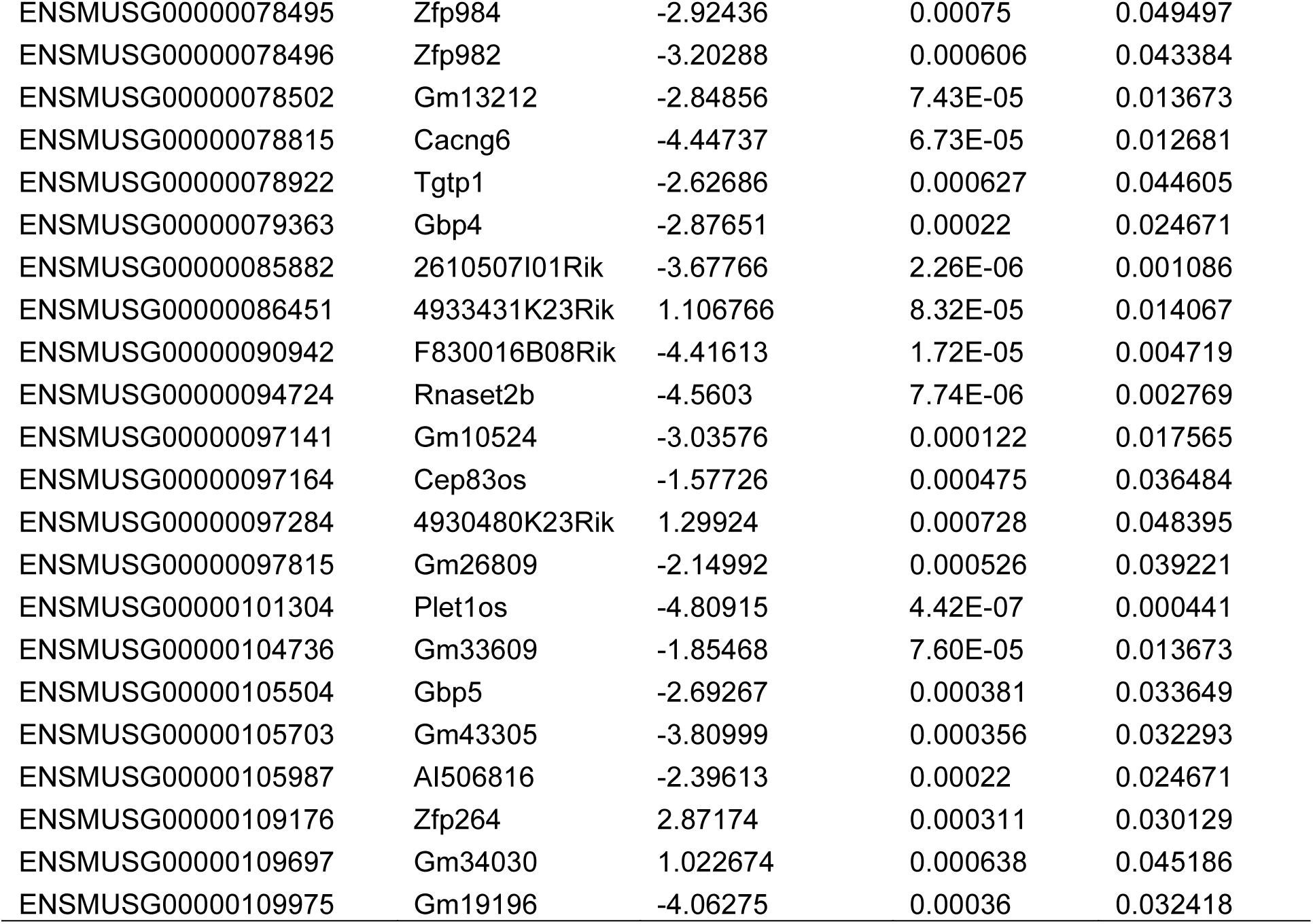
Cardiac genes significantly altered in *Il10*^-/-^ mice.

**Table S7.**
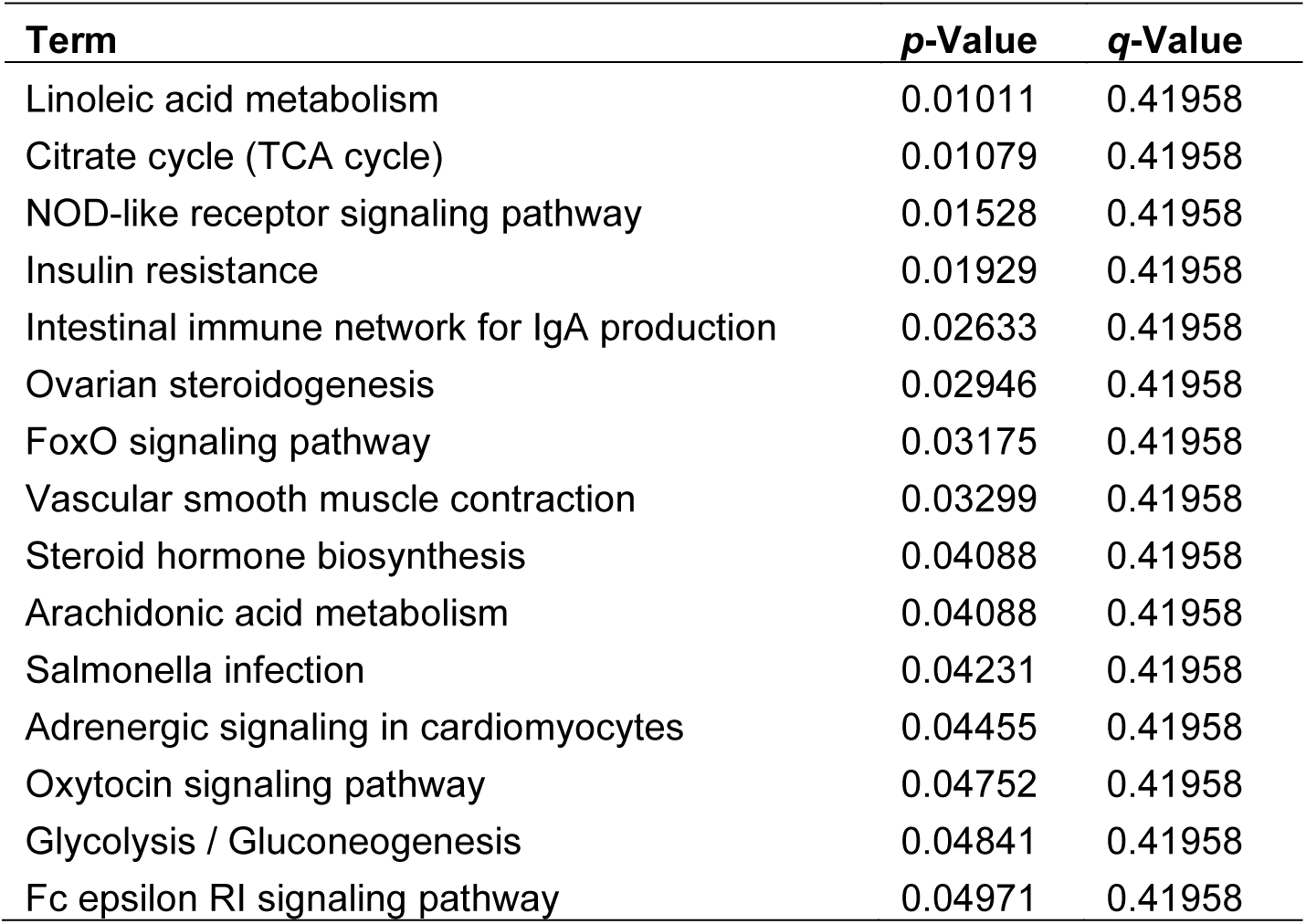
All significantly enriched signaling pathways identified by KEGG enrichment analysis (all DEGs, *Il10*^-/-^ *vs*. WT).

**Table S8.**
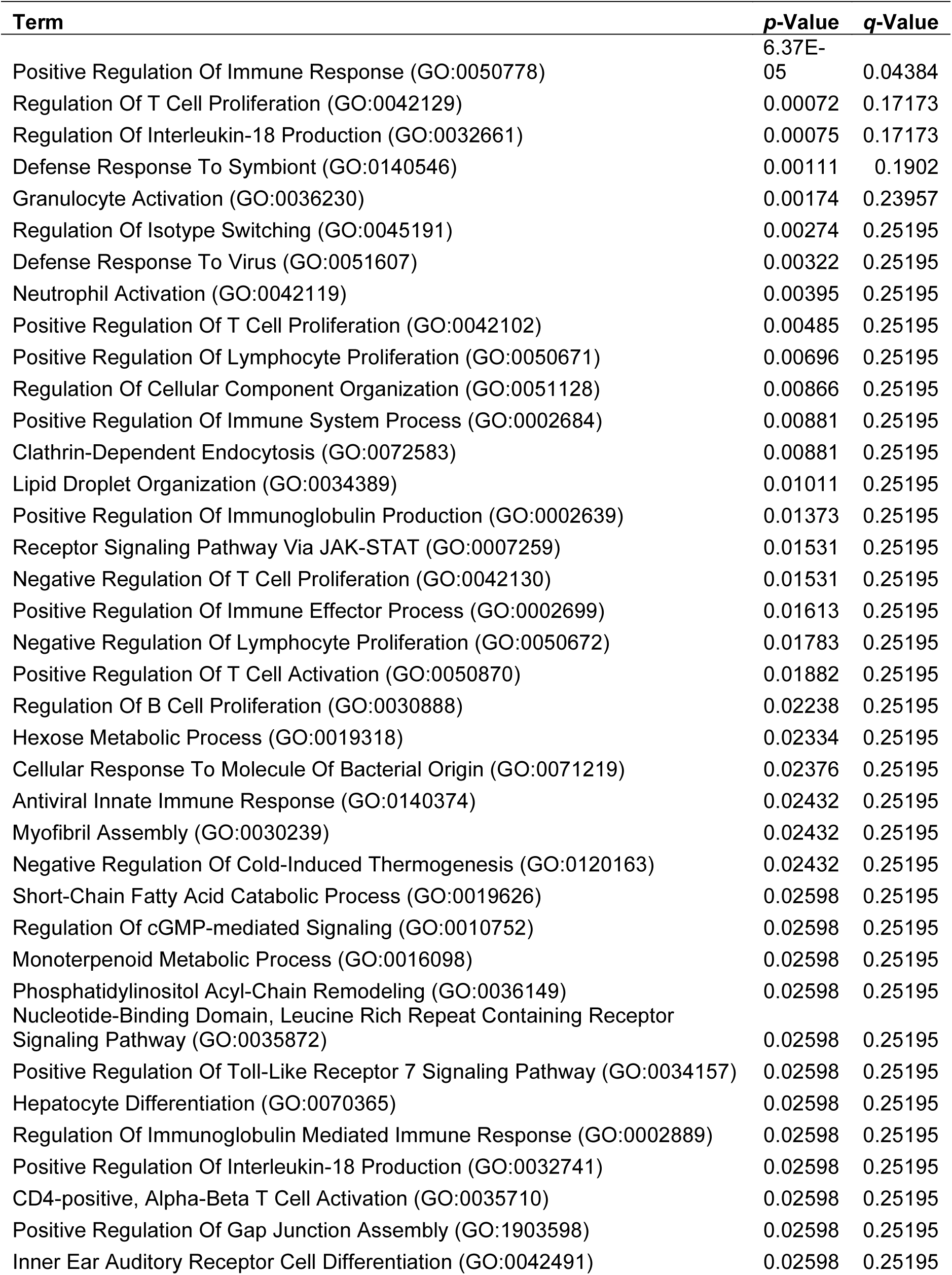
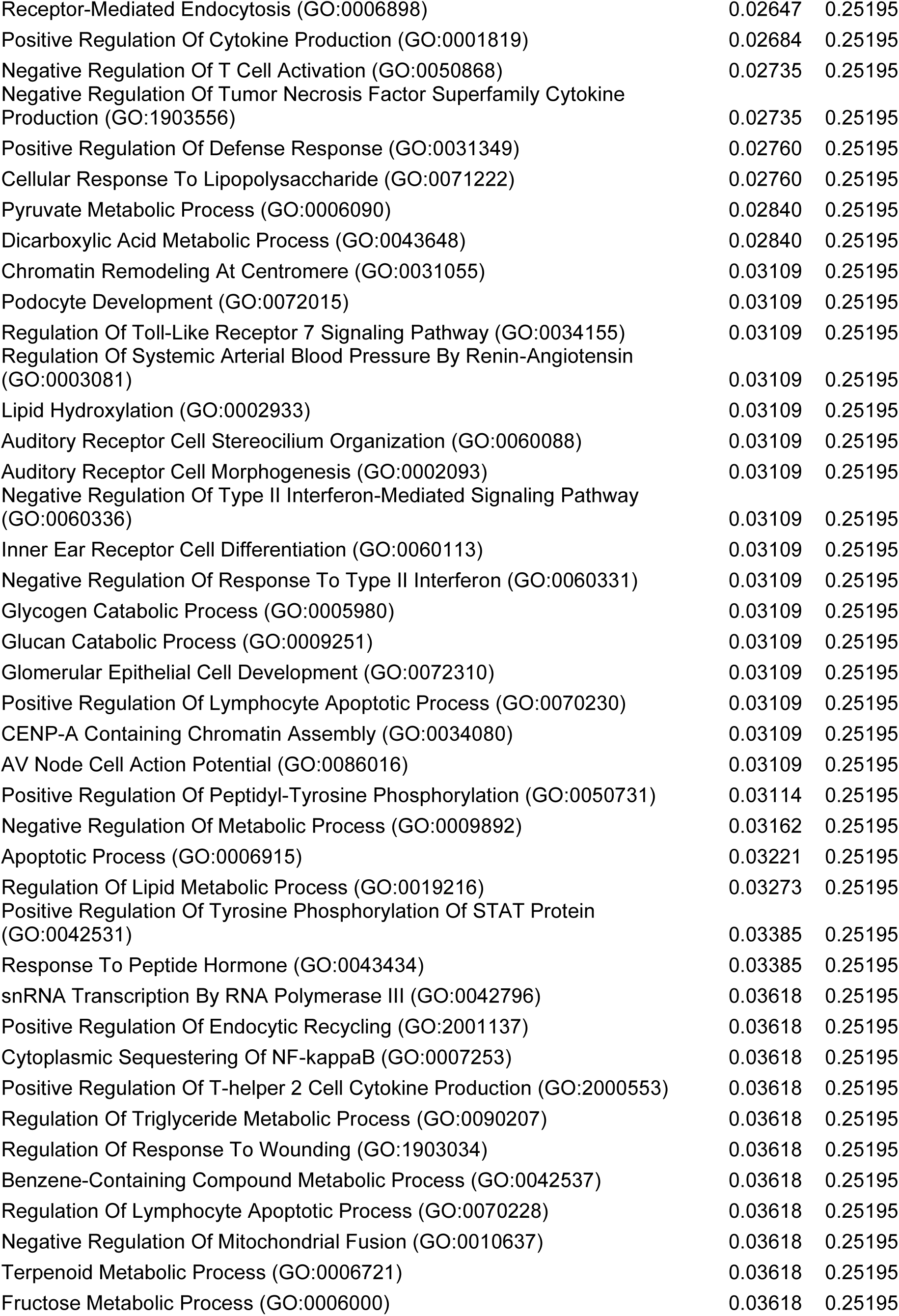
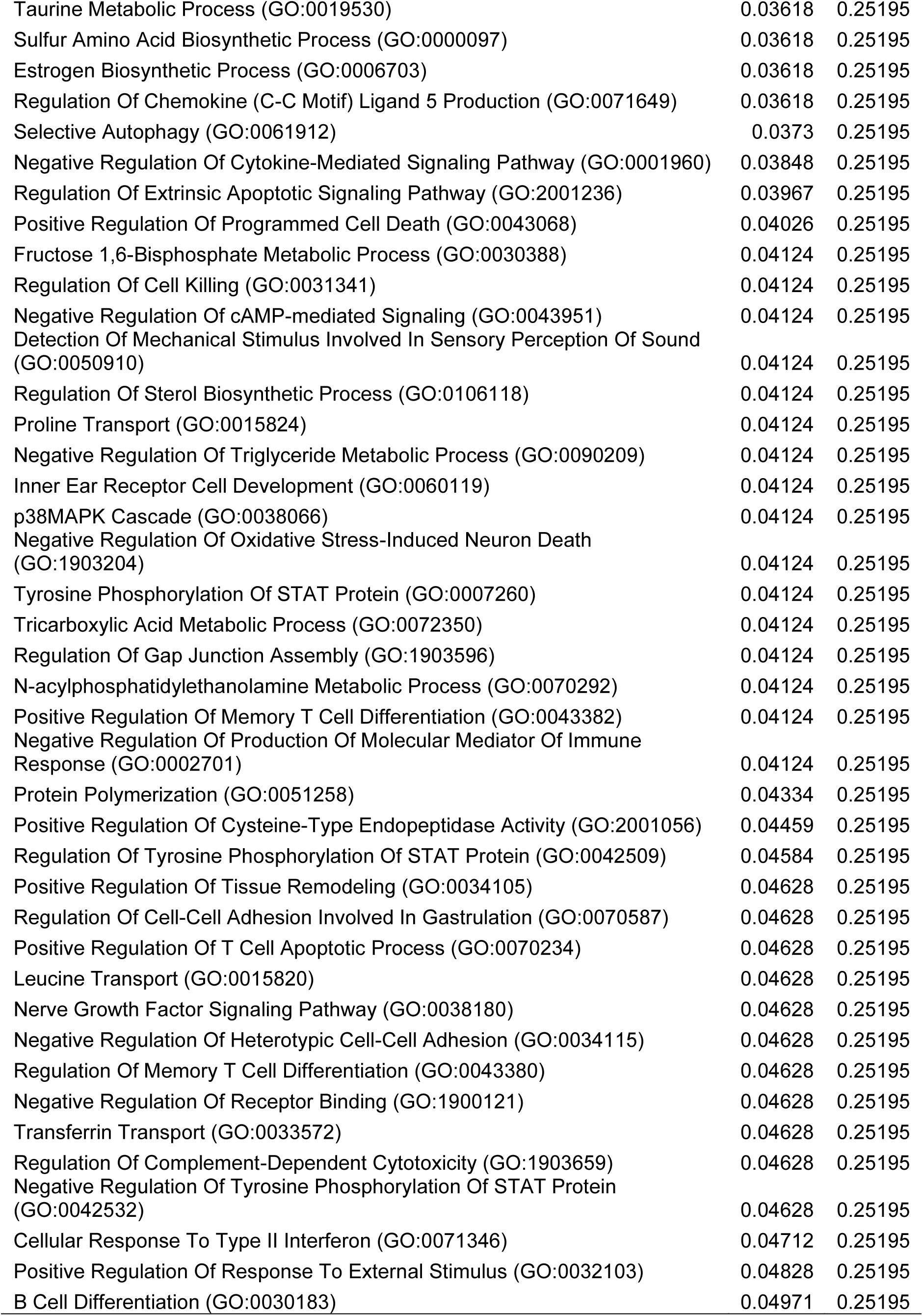
Significantly enriched biological processes identified by GO enrichment analysis (all DEGs, *Il10*^-/-^ vs. WT).

**Table S9.** Cardiac genes significantly altered by fecal microbiota transplantation in *Il10*^-/-^ mice.

| ID | Gene name | log2FoldChange | p-Value | adj. p-Value |
| --- | --- | --- | --- | --- |
| ENSMUSG00000010175 | Prox1 | 1.13655 | 5.08E-18 | 6.19E-14 |
| ENSMUSG00000015882 | Lcorl | 1.223703 | 0.000215 | 0.007944 |
| ENSMUSG00000016024 | Lbp | -1.0192 | 0.000838 | 0.017773 |
| ENSMUSG00000020108 | Ddit4 | -1.22208 | 5.24E-05 | 0.003221 |
| ENSMUSG00000020181 | Nav3 | 1.166006 | 7.17E-11 | 4.85E-08 |
| ENSMUSG00000020189 | Osbpl8 | 1.019306 | 2.72E-05 | 0.001979 |
| ENSMUSG00000020889 | Nr1d1 | -1.01603 | 0.000111 | 0.005475 |
| ENSMUSG00000021922 | Itih4 | -1.33109 | 0.000664 | 0.015809 |
| ENSMUSG00000021952 | Xpo4 | 1.352765 | 3.91E-05 | 0.002575 |
| ENSMUSG00000022868 | Ahsg | -1.04062 | 0.003164 | 0.033513 |
| ENSMUSG00000022871 | Fetub | -1.04524 | 0.000193 | 0.007428 |
| ENSMUSG00000023067 | Cdkn1a | -1.43546 | 5.32E-08 | 1.35E-05 |
| ENSMUSG00000023829 | Slc22a1 | -1.26273 | 0.001245 | 0.021773 |
| ENSMUSG00000024222 | Fkbp5 | -1.54063 | 0.002635 | 0.030785 |
| ENSMUSG00000024679 | Ms4a6d | -1.21066 | 0.000955 | 0.019035 |
| ENSMUSG00000025019 | Lcor | 1.362569 | 0.000149 | 0.006509 |
| ENSMUSG00000025938 | Slco5a1 | 1.242115 | 4.23E-05 | 0.002722 |
| ENSMUSG00000026605 | Cenpf | 1.035421 | 0.000946 | 0.018998 |
| ENSMUSG00000026822 | Lcn2 | -1.29882 | 0.000238 | 0.008379 |
| ENSMUSG00000026971 | Itgb6 | 1.026256 | 1.61E-11 | 1.51E-08 |
| ENSMUSG00000027239 | Mdk | -1.09525 | 0.002552 | 0.030214 |
| ENSMUSG00000028862 | Map3k6 | -1.88277 | 0.000657 | 0.015774 |
| ENSMUSG00000029321 | Slc10a6 | -1.18845 | 0.001798 | 0.025773 |
| ENSMUSG00000029366 | Dck | 1.052613 | 5.12E-06 | 0.000615 |
| ENSMUSG00000029675 | Eln | -1.07944 | 0.000172 | 0.00702 |
| ENSMUSG00000029727 | Cyp3a13 | 1.078672 | 0.000689 | 0.016005 |
| ENSMUSG00000030717 | Nupr1 | -1.04264 | 0.000589 | 0.014762 |
| ENSMUSG00000031558 | Slit2 | 1.184631 | 0.001836 | 0.025949 |
| ENSMUSG00000031722 | Hp | -1.33594 | 0.000146 | 0.006471 |
| ENSMUSG00000031762 | Mt2 | -2.46758 | 0.001003 | 0.019461 |
| ENSMUSG00000031765 | Mt1 | -1.20144 | 7.01E-06 | 0.000756 |
| ENSMUSG00000031994 | Adamts8 | -1.50838 | 0.003029 | 0.032856 |
| ENSMUSG00000034218 | Atm | 1.096634 | 0.003205 | 0.033718 |
| ENSMUSG00000035236 | Scai | 1.000819 | 0.002979 | 0.032593 |
| ENSMUSG00000036019 | Tmtc2 | 1.061487 | 1.16E-07 | 2.66E-05 |
| ENSMUSG00000038550 | Ciart | -1.04137 | 1.02E-09 | 5.40E-07 |
| ENSMUSG00000039084 | Chad | -1.65856 | 0.003833 | 0.037195 |
| ENSMUSG00000039956 | Mrap | -1.0994 | 0.000214 | 0.007942 |
| ENSMUSG00000040710 | St8sia4 | 1.240315 | 7.08E-08 | 1.72E-05 |
| ENSMUSG00000040855 | Reps2 | 1.164053 | 5.13E-07 | 9.91E-05 |
| ENSMUSG00000041633 | Kctd12b | 1.083528 | 8.56E-14 | 1.74E-10 |
| ENSMUSG00000042436 | Mfap4 | -1.28089 | 0.002736 | 0.031329 |
| ENSMUSG00000043467 | Zbtb37 | 1.071741 | 0.000298 | 0.009656 |
| ENSMUSG00000043613 | Mmp3 | -1.68805 | 1.17E-05 | 0.001056 |
| ENSMUSG00000044499 | Hs3st5 | 1.542304 | 1.24E-09 | 6.28E-07 |
| ENSMUSG00000044749 | Abca6 | 1.252182 | 6.13E-06 | 0.000691 |
| ENSMUSG00000046694 | Fam46b | -1.08516 | 0.001803 | 0.02581 |
| ENSMUSG00000047146 | Tet1 | 1.179904 | 3.43E-05 | 0.002335 |
| ENSMUSG00000047496 | Rnf152 | 1.052783 | 0.003103 | 0.033369 |
| ENSMUSG00000047996 | Prrg1 | 1.124163 | 3.82E-06 | 0.00048 |
| ENSMUSG00000049303 | Syt12 | -1.32908 | 0.001504 | 0.02394 |
| ENSMUSG00000052302 | Tbc1d30 | 1.042687 | 1.40E-05 | 0.001209 |
| ENSMUSG00000053205 | Styx | 1.140744 | 0.000106 | 0.005272 |
| ENSMUSG00000054659 | Pm20d2 | 1.363711 | 0.003576 | 0.036068 |
| ENSMUSG00000055639 | Dach1 | 1.246298 | 1.04E-12 | 1.27E-09 |
| ENSMUSG00000059430 | Actg2 | -1.16275 | 0.00129 | 0.022233 |
| ENSMUSG00000061718 | Ppp1r1b | -2.66097 | 0.000352 | 0.010741 |
| ENSMUSG00000061878 | Sphk1 | -1.52111 | 0.000275 | 0.009114 |
| ENSMUSG00000063108 | Zfp26 | 1.069218 | 0.00417 | 0.038747 |
| ENSMUSG00000063286 | Gm8995 | 1.04713 | 2.96E-08 | 8.38E-06 |
| ENSMUSG00000063415 | Cyp26b1 | 1.138716 | 0.00141 | 0.023201 |
| ENSMUSG00000064246 | Chil1 | -2.4146 | 0.002137 | 0.028071 |
| ENSMUSG00000067818 | Myl9 | -1.1663 | 0.002884 | 0.032024 |
| ENSMUSG00000073016 | Uprt | 1.342352 | 0.004664 | 0.041769 |
| ENSMUSG00000073664 | Nbeal1 | 1.001143 | 0.000753 | 0.016921 |
| ENSMUSG00000091971 | Hspa1a | -1.13636 | 0.000739 | 0.016751 |
| ENSMUSG00000094410 | Gm38394 | 1.719623 | 0.000417 | 0.011971 |

**Table S10.** Top 10 enriched KEGG pathways (all DEGs, *Il10*^-/-^+FMT vs *Il10*^-/-^).

| Term | p-Value | q-Value | Overlap genes |
| --- | --- | --- | --- |
| Transcriptional misregulation in cancer | 0.003938 | 0.385965 | [CDKN1A, MMP3, ATM, NUPR1] |
| p53 signaling pathway | 0.024939 | 0.457175 | [CDKN1A, ATM] |
| Human papillomavirus infection | 0.024989 | 0.457175 | [CDKN1A, CHAD, ATM, ITGB6] |
| Focal adhesion | 0.029949 | 0.457175 | [CHAD, ITGB6, MYL9] |
| PI3K-Akt signaling pathway | 0.030932 | 0.457175 | [CDKN1A, DDIT4, CHAD, ITGB6] |
| ECM-receptor interaction | 0.035185 | 0.457175 | [CHAD, ITGB6] |
| Lipid and atherosclerosis | 0.035508 | 0.457175 | [MMP3, LBP, HSPA1A] |
| IL-17 signaling pathway | 0.039667 | 0.457175 | [MMP3, LCN2] |
| Prostate cancer | 0.041985 | 0.457175 | [CDKN1A, MMP3] |
| NF-kappa B signaling pathway | 0.047587 | 0.461354 | [ATM, LBP] |

**Table S11.** Significantly enriched biological processes identified by GO enrichment analysis (all DEGs, *Il10*^-/-+^FMT vs. *Il10*^-/-^).

| Term | p-value | q-value |
| --- | --- | --- |
| Regulation Of Neuroinflammatory Response (GO:0150077) | 4.29E-07 | 0.00034 |
| Negative Regulation Of Cell Motility (GO:2000146) | 8.37E-05 | 0.02194 |
| Regulation Of Cell Migration (GO:0030334) | 9.81E-05 | 0.02194 |
| Negative Regulation Of Cardiac Muscle Cell Apoptotic Process (GO:0010667) | 0.00011 | 0.02194 |
| Regulation Of DNA Biosynthetic Process (GO:2000278) | 0.00020 | 0.02621 |
| Negative Regulation Of Cell Migration (GO:0030336) | 0.00022 | 0.02621 |
| Negative Regulation Of Striated Muscle Cell Apoptotic Process (GO:0010664) | 0.00023 | 0.02621 |
| Regulation Of Bile Acid Biosynthetic Process (GO:0070857) | 0.000392 | 0.03915 |
| Regulation Of Cardiac Muscle Cell Apoptotic Process (GO:0010665) | 0.000489 | 0.04341 |
| Regulation Of Epithelial Cell Apoptotic Process (GO:1904035) | 0.000596 | 0.04764 |
| Negative Regulation Of TOR Signaling (GO:0032007) | 0.000789 | 0.05729 |
| Regulation Of Microglial Cell Activation (GO:1903978) | 0.00098 | 0.06527 |
| Regulation Of Inflammatory Response (GO:0050727) | 0.00126 | 0.07268 |
| Positive Regulation Of Cell Cycle (GO:0045787) | 0.00135 | 0.07268 |
| Regulation Of Leukocyte Chemotaxis (GO:0002688) | 0.00146 | 0.07268 |
| Extracellular Matrix Assembly (GO:0085029) | 0.00146 | 0.07268 |
| Regulation Of Cell Cycle G2/M Phase Transition (GO:1902749) | 0.00163 | 0.07679 |
| Negative Regulation Of Cellular Process (GO:0048523) | 0.00209 | 0.09284 |
| Cellular Response To UV (GO:0034644) | 0.00237 | 0.09305 |
| Regulation Of Mononuclear Cell Migration (GO:0071675) | 0.00245 | 0.09305 |
| Regulation Of Neural Precursor Cell Proliferation (GO:2000177) | 0.00245 | 0.09305 |
| Regulation Of Neutrophil Chemotaxis (GO:0090022) | 0.00316 | 0.10095 |
| Mitotic Spindle Checkpoint Signaling (GO:0071174) | 0.00341 | 0.10095 |
| Mitotic Spindle Assembly Checkpoint Signaling (GO:0007094) | 0.00341 | 0.10095 |
| Spindle Assembly Checkpoint Signaling (GO:0071173) | 0.00341 | 0.10095 |
| Acute Inflammatory Response (GO:0002526) | 0.00341 | 0.10095 |
| Protein Export From Nucleus (GO:0006611) | 0.00341 | 0.10095 |
| Ventricular Septum Morphogenesis (GO:0060412) | 0.00368 | 0.10251 |
| Negative Regulation Of Mitotic Metaphase/Anaphase Transition (GO:0045841) | 0.00395 | 0.10251 |
| Positive Regulation Of Fibroblast Proliferation (GO:0048146) | 0.00395 | 0.10251 |
| Negative Regulation Of Plasma Membrane Bounded Cell Projection Assembly (GO:0120033) | 0.00423 | 0.10251 |
| Positive Regulation Of Microtubule Polymerization (GO:0031116) | 0.00423 | 0.10251 |
| Positive Regulation Of Neural Precursor Cell Proliferation (GO:2000179) | 0.00423 | 0.10251 |
| Mitotic G2 DNA Damage Checkpoint Signaling (GO:0007095) | 0.00453 | 0.10637 |
| Negative Regulation Of DNA Biosynthetic Process (GO:2000279) | 0.00483 | 0.11022 |
| Lipid Transport (GO:0006869) | 0.00571 | 0.12501 |
| Regulation Of G2/M Transition Of Mitotic Cell Cycle (GO:0010389) | 0.00613 | 0.12501 |
| DNA Damage Response, Signal Transduction By P53 Class Mediator (GO:0030330) | 0.00647 | 0.12501 |
| Negative Regulation Of Organelle Assembly (GO:1902116) | 0.00647 | 0.12501 |
| Aortic Valve Morphogenesis (GO:0003180) | 0.00683 | 0.12501 |
| Regulation Of Ossification (GO:0030278) | 0.00683 | 0.12501 |
| Ventricular Septum Development (GO:0003281) | 0.00683 | 0.12501 |
| Negative Regulation Of DNA-templated Transcription (GO:0045892) | 0.00695 | 0.12501 |
| Regulation Of Cell Growth (GO:0001558) | 0.00709 | 0.12501 |
| Negative Regulation Of Epithelial Cell Apoptotic Process (GO:1904036) | 0.00719 | 0.12501 |
| Cellular Response To Lipid (GO:0071396) | 0.00720 | 0.12501 |
| Aortic Valve Development (GO:0003176) | 0.00794 | 0.13504 |
| Response To Amino Acid Starvation (GO:1990928) | 0.00833 | 0.13517 |
| Negative Regulation Of Growth (GO:0045926) | 0.00835 | 0.13517 |
| Negative Regulation Of Cell Growth (GO:0030308) | 0.00854 | 0.13517 |
| Cellular Response To Amino Acid Starvation (GO:0034198) | 0.00873 | 0.13517 |
| Negative Regulation Of Cell Population Proliferation (GO:0008285) | 0.00880 | 0.13517 |
| Regulation Of Primary Metabolic Process (GO:0080090) | 0.00914 | 0.13775 |
| Organic Hydroxy Compound Transport (GO:0015850) | 0.00997 | 0.14227 |
| Regulation Of Fibroblast Proliferation (GO:0048145) | 0.01041 | 0.14227 |
| Mitotic G2/M Transition Checkpoint (GO:0044818) | 0.01041 | 0.14227 |
| Negative Regulation Of Cellular Biosynthetic Process (GO:0031327) | 0.01095 | 0.14227 |
| Regulation Of Tumor Necrosis Factor-Mediated Signaling Pathway (GO:0010803) | 0.01129 | 0.14227 |
| Cellular Response To Oxygen-Containing Compound (GO:1901701) | 0.01161 | 0.14227 |
| Intrinsic Apoptotic Signaling Pathway In Response To DNA Damage (GO:0008630) | 0.01175 | 0.14227 |
| Negative Regulation Of DNA Metabolic Process (GO:0051053) | 0.01175 | 0.14227 |
| Positive Regulation Of Cell Death (GO:0010942) | 0.01221 | 0.14227 |
| Regulation Of TORC1 Signaling (GO:1903432) | 0.01221 | 0.14227 |
| Negative Regulation Of Transcription By RNA Polymerase II (GO:0000122) | 0.01369 | 0.14227 |
| Positive Regulation Of Cellular Process (GO:0048522) | 0.01445 | 0.14227 |
| Mitotic DNA Damage Checkpoint Signaling (GO:0044773) | 0.01515 | 0.14227 |
| Regulation Of Phagocytosis (GO:0050764) | 0.01620 | 0.14227 |
| Renal System Development (GO:0072001) | 0.01620 | 0.14227 |
| Negative Regulation Of Chemokine-Mediated Signaling Pathway (GO:0070100) | 0.01664 | 0.14227 |
| Negative Regulation Of Astrocyte Differentiation (GO:0048712) | 0.01664 | 0.14227 |
| Leukocyte Chemotaxis Involved In Inflammatory Response (GO:0002232) | 0.01664 | 0.14227 |
| Iron Coordination Entity Transport (GO:1901678) | 0.01664 | 0.14227 |
| Xenobiotic Transport Across Blood-Brain Barrier (GO:1990962) | 0.01664 | 0.14227 |
| Hepatocyte Differentiation (GO:0070365) | 0.01664 | 0.14227 |
| Heat Acclimation (GO:0010286) | 0.01664 | 0.14227 |
| Ventricular Cardiac Muscle Cell Development (GO:0055015) | 0.01664 | 0.14227 |
| Venous Blood Vessel Morphogenesis (GO:0048845) | 0.01664 | 0.14227 |
| Thiamine Transmembrane Transport (GO:0071934) | 0.01664 | 0.14227 |
| Retinal Ganglion Cell Axon Guidance (GO:0031290) | 0.01664 | 0.14227 |
| Response To Heparin (GO:0071503) | 0.01664 | 0.14227 |
| Negative Regulation Of Monocyte Chemotaxis (GO:0090027) | 0.01664 | 0.14227 |
| Polyamine Transport (GO:0015846) | 0.01664 | 0.14227 |
| Regulation Of Sarcomere Organization (GO:0060297) | 0.01664 | 0.14227 |
| Cellular Heat Acclimation (GO:0070370) | 0.01664 | 0.14227 |
| Regulation Of Retinal Ganglion Cell Axon Guidance (GO:0090259) | 0.01664 | 0.14227 |
| Regulation Of Neutrophil Extravasation (GO:2000389) | 0.01664 | 0.14227 |
| Negative Regulation Of Sequestering Of Triglyceride (GO:0010891) | 0.01664 | 0.14227 |
| Regulation Of Hydrogen Peroxide Metabolic Process (GO:0010310) | 0.01664 | 0.14227 |
| Regulation Of Hepatocyte Proliferation (GO:2000345) | 0.01664 | 0.14227 |
| Negative Regulation Of Lamellipodium Organization (GO:1902744) | 0.01664 | 0.14227 |
| Negative Regulation Of Heterochromatin Formation (GO:0031452) | 0.01664 | 0.14227 |
| Regulation Of Circadian Sleep/Wake Cycle (GO:0042749) | 0.01664 | 0.14227 |
| Regulation Of Type B Pancreatic Cell Proliferation (GO:0061469) | 0.01664 | 0.14227 |
| Regulation Of Cell Cycle (GO:0051726) | 0.01777 | 0.14227 |
| Negative Regulation Of Nucleic Acid-Templated Transcription (GO:1903507) | 0.01832 | 0.14227 |
| Regulation Of Cellular Catabolic Process (GO:0031329) | 0.01893 | 0.14227 |
| Brain Development (GO:0007420) | 0.01913 | 0.14227 |
| Negative Regulation Of Retinoic Acid Receptor Signaling Pathway (GO:0048387) | 0.01993 | 0.14227 |
| Neuron Projection Extension Involved In Neuron Projection Guidance (GO:1902284) | 0.01993 | 0.14227 |
| Neurotrophin TRK Receptor Signaling Pathway (GO:0048011) | 0.01993 | 0.14227 |
| Axon Extension Involved In Axon Guidance (GO:0048846) | 0.01993 | 0.14227 |
| Regulation Of Nucleotide-Binding Oligomerization Domain Containing 2 Signaling Pathway (GO:0070432) | 0.01993 | 0.14227 |
| Nose Development (GO:0043584) | 0.01993 | 0.14227 |
| Catecholamine Uptake (GO:0090493) | 0.01993 | 0.14227 |
| Regulation Of Striated Muscle Tissue Development (GO:0016202) | 0.01993 | 0.14227 |
| Elastic Fiber Assembly (GO:0048251) | 0.01993 | 0.14227 |
| Skeletal Muscle Thin Filament Assembly (GO:0030240) | 0.01993 | 0.14227 |
| Response To Reactive Oxygen Species (GO:0000302) | 0.02066 | 0.14227 |
| Negative Regulation Of Cellular Catabolic Process (GO:0031330) | 0.02125 | 0.14227 |
| Extracellular Matrix Organization (GO:0030198) | 0.02127 | 0.14227 |
| Regulation Of Endopeptidase Activity (GO:0052548) | 0.02245 | 0.14227 |
| Lymphatic Endothelial Cell Differentiation (GO:0060836) | 0.02322 | 0.14227 |
| Lens Fiber Cell Development (GO:0070307) | 0.02322 | 0.14227 |
| Positive Regulation Of Steroid Biosynthetic Process (GO:0010893) | 0.02322 | 0.14227 |
| Positive Regulation Of Oligodendrocyte Differentiation (GO:0048714) | 0.02322 | 0.14227 |
| Venous Blood Vessel Development (GO:0060841) | 0.02322 | 0.14227 |
| Positive Regulation Of Microtubule Nucleation (GO:0090063) | 0.02322 | 0.14227 |
| Stress-Induced Premature Senescence (GO:0090400) | 0.02322 | 0.14227 |
| Dopamine Uptake (GO:0090494) | 0.02322 | 0.14227 |
| Response To Auditory Stimulus (GO:0010996) | 0.02322 | 0.14227 |
| Cellular Response To Reactive Nitrogen Species (GO:1902170) | 0.02322 | 0.14227 |
| Cellular Response To Nitric Oxide (GO:0071732) | 0.02322 | 0.14227 |
| Regulation Of Transforming Growth Factor Beta Activation (GO:1901388) | 0.02322 | 0.14227 |
| Phosphatidylserine Acyl-Chain Remodeling (GO:0036150) | 0.02322 | 0.14227 |
| Catecholamine Transport (GO:0051937) | 0.02322 | 0.14227 |
| Regulation Of Oxidative Stress-Induced Cell Death (GO:1903201) | 0.02322 | 0.14227 |
| UV Protection (GO:0009650) | 0.02322 | 0.14227 |
| Negative Regulation Of Microglial Cell Activation (GO:1903979) | 0.02322 | 0.14227 |
| Negative Regulation Of Inflammatory Response To Wounding (GO:0106015) | 0.02322 | 0.14227 |
| Regulation Of Chemokine-Mediated Signaling Pathway (GO:0070099) | 0.02322 | 0.14227 |
| Negative Regulation Of Supramolecular Fiber Organization (GO:1902904) | 0.02368 | 0.14227 |
| Kidney Development (GO:0001822) | 0.02368 | 0.14227 |
| Negative Regulation Of Protein Binding (GO:0032091) | 0.02368 | 0.14227 |
| Response To Ionizing Radiation (GO:0010212) | 0.02431 | 0.14409 |
| Gland Development (GO:0048732) | 0.02558 | 0.14409 |
| Negative Regulation Of Phosphorylation (GO:0042326) | 0.02622 | 0.14409 |
| Pyrimidine-Containing Compound Metabolic Process (GO:0072527) | 0.02649 | 0.14409 |
| Neurotransmitter Reuptake (GO:0098810) | 0.02649 | 0.14409 |
| Regulation Of Vesicle Fusion (GO:0031338) | 0.02649 | 0.14409 |
| Response To Leptin (GO:0044321) | 0.02649 | 0.14409 |
| Serotonin Uptake (GO:0051610) | 0.02649 | 0.14409 |
| Skeletal Myofibril Assembly (GO:0014866) | 0.02649 | 0.14409 |
| Fat-Soluble Vitamin Catabolic Process (GO:0042363) | 0.02649 | 0.14409 |
| Positive Regulation Of Tumor Necrosis Factor-Mediated Signaling Pathway (GO:1903265) | 0.02649 | 0.14409 |
| Positive Regulation Of Endothelial Cell Proliferation (GO:0001938) | 0.02753 | 0.14409 |
| Negative Regulation Of Transforming Growth Factor Beta Receptor Signaling Pathway (GO:0030512) | 0.02753 | 0.14409 |
| Negative Regulation Of Protein Modification Process (GO:0031400) | 0.02820 | 0.14409 |
| Regulation Of Cyclin-Dependent Protein Serine/Threonine Kinase Activity (GO:0000079) | 0.02887 | 0.14409 |
| Motor Neuron Axon Guidance (GO:0008045) | 0.02975 | 0.14409 |
| Kinetochore Assembly (GO:0051382) | 0.02975 | 0.14409 |
| Positive Regulation Of Smooth Muscle Contraction (GO:0045987) | 0.02975 | 0.14409 |
| Glycogen Biosynthetic Process (GO:0005978) | 0.02975 | 0.14409 |
| Glucan Biosynthetic Process (GO:0009250) | 0.02975 | 0.14409 |
| Smooth Muscle Tissue Development (GO:0048745) | 0.02975 | 0.14409 |
| Serotonin Transport (GO:0006837) | 0.02975 | 0.14409 |
| Dopamine Transport (GO:0015872) | 0.02975 | 0.14409 |
| Toxin Transport (GO:1901998) | 0.02975 | 0.14409 |
| Autophagy Of Peroxisome (GO:0030242) | 0.02975 | 0.14409 |
| Negative Regulation Of Transcription By Competitive Promoter Binding (GO:0010944) | 0.02975 | 0.14409 |
| Nucleobase-Containing Compound Biosynthetic Process (GO:0034654) | 0.02975 | 0.14409 |
| Negative Regulation Of Toll-Like Receptor 4 Signaling Pathway (GO:0034144) | 0.02975 | 0.14409 |
| Adrenal Gland Development (GO:0030325) | 0.02975 | 0.14409 |
| Regulation Of Inflammatory Response To Wounding (GO:0106014) | 0.02975 | 0.14409 |
| Negative Regulation Of Leukocyte Migration (GO:0002686) | 0.02975 | 0.14409 |
| Regulation Of Ceramide Biosynthetic Process (GO:2000303) | 0.02975 | 0.14409 |
| Organic Substance Transport (GO:0071702) | 0.02995 | 0.14415 |
| Negative Regulation Of Autophagosome Assembly (GO:1902902) | 0.03301 | 0.14418 |
| Monoamine Transport (GO:0015844) | 0.03301 | 0.14418 |
| Prostaglandin Transport (GO:0015732) | 0.03301 | 0.14418 |
| Kinetochore Organization (GO:0051383) | 0.03301 | 0.14418 |
| Diol Metabolic Process (GO:0034311) | 0.03301 | 0.14418 |
| Positive Regulation Of Cellular Respiration (GO:1901857) | 0.03301 | 0.14418 |
| Cellular Response To UV-A (GO:0071492) | 0.03301 | 0.14418 |
| Regulation Of Programmed Necrotic Cell Death (GO:0062098) | 0.03301 | 0.14418 |
| Negative Regulation Of Oxidoreductase Activity (GO:0051354) | 0.03301 | 0.14418 |
| Negative Regulation Of Steroid Biosynthetic Process (GO:0010894) | 0.03301 | 0.14418 |
| Negative Regulation Of Programmed Necrotic Cell Death (GO:0062099) | 0.03301 | 0.14418 |
| Regulation Of Inclusion Body Assembly (GO:0090083) | 0.03301 | 0.14418 |
| Negative Regulation Of Mitochondrial Outer Membrane Permeabilization Involved In Apoptotic Signaling Pathway (GO:1901029) | 0.03301 | 0.14418 |
| Negative Regulation Of Inclusion Body Assembly (GO:0090084) | 0.03301 | 0.14418 |
| Negative Regulation Of Glycolytic Process (GO:0045820) | 0.03301 | 0.14418 |
| Regulation Of Microtubule Nucleation (GO:0010968) | 0.03301 | 0.14418 |
| Negative Regulation Of Cell Cycle (GO:0045786) | 0.03302 | 0.14418 |
| Negative Regulation Of Binding (GO:0051100) | 0.03374 | 0.14474 |
| Regulation Of Mitotic Cell Cycle Phase Transition (GO:1901990) | 0.03374 | 0.14474 |
| Regulation Of Endothelial Cell Migration (GO:0010594) | 0.03518 | 0.14474 |
| Positive Regulation Of Cell Adhesion (GO:0045785) | 0.03518 | 0.14474 |
| Cellular Response To Chemical Stress (GO:0062197) | 0.03592 | 0.14474 |
| Positive Regulation Of Muscle Contraction (GO:0045933) | 0.03625 | 0.14474 |
| Sphingoid Metabolic Process (GO:0046519) | 0.03625 | 0.14474 |
| Response To Nitric Oxide (GO:0071731) | 0.03625 | 0.14474 |
| Response To X-ray (GO:0010165) | 0.03625 | 0.14474 |
| Cellular Response To UV-B (GO:0071493) | 0.03625 | 0.14474 |
| Positive Regulation Of Keratinocyte Proliferation (GO:0010838) | 0.03625 | 0.14474 |
| Nucleoside Phosphate Biosynthetic Process (GO:1901293) | 0.03625 | 0.14474 |
| Regulation Of Muscle System Process (GO:0090257) | 0.03625 | 0.14474 |
| Negative Regulation Of Vascular Permeability (GO:0043116) | 0.03625 | 0.14474 |
| Regulation Of Leukocyte Cell-Cell Adhesion (GO:1903037) | 0.03625 | 0.14474 |
| Cell Differentiation In Hindbrain (GO:0021533) | 0.03625 | 0.14474 |
| Negative Regulation Of Leukocyte Chemotaxis (GO:0002689) | 0.03625 | 0.14474 |
| Positive Regulation Of Proteasomal Protein Catabolic Process (GO:1901800) | 0.03666 | 0.14474 |
| Regulation Of Epithelial Cell Proliferation (GO:0050678) | 0.03740 | 0.14474 |
| Positive Regulation Of Cell Motility (GO:2000147) | 0.03804 | 0.14474 |
| Protein O-linked Glycosylation (GO:0006493) | 0.03891 | 0.14474 |
| Positive Regulation Of Protein Export From Nucleus (GO:0046827) | 0.03948 | 0.14474 |
| Positive Regulation Of Neuron Migration (GO:2001224) | 0.03948 | 0.14474 |
| Positive Regulation Of Neuroinflammatory Response (GO:0150078) | 0.03948 | 0.14474 |
| Folic Acid Metabolic Process (GO:0046655) | 0.03948 | 0.14474 |
| Positive Regulation Of Cellular Extravasation (GO:0002693) | 0.03948 | 0.14474 |
| Negative Regulation Of Mononuclear Cell Migration (GO:0071676) | 0.03948 | 0.14474 |
| Nucleotide Metabolic Process (GO:0009117) | 0.03948 | 0.14474 |
| Atrial Cardiac Muscle Tissue Development (GO:0003228) | 0.03948 | 0.14474 |
| Regulation Of Leukocyte Adhesion To Vascular Endothelial Cell (GO:1904994) | 0.03948 | 0.14474 |
| Positive Regulation Of Vascular Endothelial Cell Proliferation (GO:1905564) | 0.03948 | 0.14474 |
| Negative Regulation Of Mitochondrial Membrane Permeability (GO:0035795) | 0.03948 | 0.14474 |
| Regulation Of Retinoic Acid Receptor Signaling Pathway (GO:0048385) | 0.03948 | 0.14474 |
| Quaternary Ammonium Group Transport (GO:0015697) | 0.03948 | 0.14474 |
| Cellular Response To Hormone Stimulus (GO:0032870) | 0.03967 | 0.14474 |
| Positive Regulation Of Supramolecular Fiber Organization (GO:1902905) | 0.04121 | 0.14474 |
| Negative Regulation Of Intracellular Signal Transduction (GO:1902532) | 0.04247 | 0.14474 |
| Negative Regulation Of Multicellular Organismal Process (GO:0051241) | 0.04247 | 0.14474 |
| Negative Regulation Of Cell Activation (GO:0050866) | 0.04270 | 0.14474 |
| Negative Regulation Of T Cell Differentiation (GO:0045581) | 0.04270 | 0.14474 |
| Positive Regulation Of Macrophage Chemotaxis (GO:0010759) | 0.04270 | 0.14474 |
| Sphingosine Biosynthetic Process (GO:0046512) | 0.04270 | 0.14474 |
| Sodium-Independent Organic Anion Transport (GO:0043252) | 0.04270 | 0.14474 |
| Endothelial Cell Differentiation (GO:0045446) | 0.04270 | 0.14474 |
| Sphingoid Biosynthetic Process (GO:0046520) | 0.04270 | 0.14474 |
| Regulation Of Response To Endoplasmic Reticulum Stress (GO:1905897) | 0.04270 | 0.14474 |
| Regulation Of Oxidoreductase Activity (GO:0051341) | 0.04270 | 0.14474 |
| Regulation Of Nitrogen Compound Metabolic Process (GO:0051171) | 0.04270 | 0.14474 |
| Acute-Phase Response (GO:0006953) | 0.04270 | 0.14474 |
| Negative Regulation Of Neuroinflammatory Response (GO:0150079) | 0.04270 | 0.14474 |
| Response To UV-A (GO:0070141) | 0.04270 | 0.14474 |
| Negative Regulation Of Macromolecule Biosynthetic Process (GO:0010558) | 0.04277 | 0.14474 |
| Positive Regulation Of Protein-Containing Complex Assembly (GO:0031334) | 0.04277 | 0.14474 |
| Regulation Of Cell Population Proliferation (GO:0042127) | 0.04293 | 0.14474 |
| Striated Muscle Cell Development (GO:0055002) | 0.04591 | 0.14850 |
| Regulation Of Bone Remodeling (GO:0046850) | 0.04591 | 0.14850 |
| Icosanoid Transport (GO:0071715) | 0.04591 | 0.14850 |
| Establishment Of Chromosome Localization (GO:0051303) | 0.04591 | 0.14850 |
| Negative Regulation Of Cilium Assembly (GO:1902018) | 0.04591 | 0.14850 |
| Regulation Of Sequestering Of Triglyceride (GO:0010889) | 0.04591 | 0.14850 |
| Pancreas Development (GO:0031016) | 0.04591 | 0.14850 |
| Bone Morphogenesis (GO:0060349) | 0.04591 | 0.14850 |
| Negative Regulation Of Smooth Muscle Cell Migration (GO:0014912) | 0.04591 | 0.14850 |
| Centromere Complex Assembly (GO:0034508) | 0.04591 | 0.14850 |
| Regulation Of Autophagy (GO:0010506) | 0.04714 | 0.15090 |
| Positive Regulation Of Inflammatory Response (GO:0050729) | 0.04759 | 0.15090 |
| Regulation Of Neuron Apoptotic Process (GO:0043523) | 0.04841 | 0.15090 |
| Intrinsic Apoptotic Signaling Pathway (GO:0097193) | 0.04841 | 0.15090 |
| Ganglioside Biosynthetic Process (GO:0001574) | 0.04911 | 0.15090 |
| Sphingolipid Catabolic Process (GO:0030149) | 0.04911 | 0.15090 |
| Positive Regulation Of Cyclin-Dependent Protein Serine/Threonine Kinase Activity (GO:0045737) | 0.04911 | 0.15090 |
| Positive Regulation Of DNA Damage Response, Signal Transduction By P53 Class Mediator (GO:0043517) | 0.04911 | 0.15090 |
| Neurotrophin Signaling Pathway (GO:0038179) | 0.04911 | 0.15090 |
| Oogenesis (GO:0048477) | 0.04911 | 0.15090 |
| Negative Regulation Of Macrophage Activation (GO:0043031) | 0.04911 | 0.15090 |
| Negative Regulation Of Endoplasmic Reticulum Stress-Induced Intrinsic Apoptotic Signaling Pathway (GO:1902236) | 0.04911 | 0.15090 |
| Cardiac Myofibril Assembly (GO:0055003) | 0.04911 | 0.15090 |

